# Atypical Cell Cycle Regulation over Neural Stem Cell Expansion

**DOI:** 10.1101/2023.12.22.572899

**Authors:** Dorota Lubanska, Ingrid Qemo, Keith Franklin Stringer, Hema Priya Mahendran, Bre-Anne Fifield, Alan Cieslukowski, Sami Alrashed, Youshaa Elabed, Emmanuel Boujeke, Alexander Rodzinka, Elizabeth Fidalgo da Silva, Stephanie Dinescu, Alexandra Sorge, Srinath Kandalam, Jillian Brown, Hasan Ghafoor, Dalton Liwanpo, Lisa A. Porter

## Abstract

Populations of adult neural stem cells (NSCs) that reside in the mammalian brain aid in neurogenesis throughout life and can be identified by a type VI intermediate filament protein, Nestin. Cell cycle regulation plays an important role in maintaining a balance between self-renewal and differentiation and determining the fate of NSCs. Data from our group and others support that the atypical cyclin-like protein Spy1 (also called RingoA; gene *SPDYA*) plays a critical role in activating NSCs from a quiescent state. Elevated levels of Spy1 are found in aggressive human brain cancers, including glioblastoma. Using a conditional mouse model, we demonstrate that driving the expression of Spy1, in the Nestin-enriched NSC population of the brain, increases stemness characteristics, decreases differentiation, and increases susceptibility to oncogenic transformation. This study contributes to better understanding of intricate cell cycle mechanisms which lead to deviation from the homeostatic state, promoting aberrant changes in adult NSCs.

## INTRODUCTION

Long-lived neural stem cells (NSC) possess the remarkable ability to self-renew and differentiate into the functional cell types within the brain. Molecular markers have been used in attempt to unambiguously categorize NSCs and their progeny in the adult SVZ. Nestin is one marker of NSCs and transient amplifying cell populations (progenitor cells) of the SVZ (Lois and Alvarez-Buylla 1994). Data suggests that Nestin is necessary for NSC self-renewal and survival (Park et al. 2010). Nestin-positive cells are exclusively found in regions of the brain that harbor a niche tailored to the maintenance of stem and progenitor cell populations (Namiki et al. 2012) where NSCs are maintained in a state of quiescence (Urbán et al. 2019). Triggering of neurogenesis in adult NSCs requires exiting quiescence and re-entering the cell cycle (Neumüller and Knoblich 2009).

Spy1 protein (gene *SPDYA*; also called RingoA) directly binds to, and activates, both CDK1 and CDK2 in a unique fashion that is independent of activating phosphorylation and dephosphorylation events (Cheng et al. 2005; McGrath et al. 2017). As a result of this mechanism Spy1 has been isolated from numerous screens and has been described for the ability to override cell cycle arrest mechanisms including DNA damage and senescence (Lenormand et al. 1999), (Lubanska et al. 2021). Work from the Nebreda lab has demonstrated that Spy1 is required for activation of NSCs out of quiescent state via activation of CDK2 and that brains of mice deficient in Spy1 have decreased neurogenesis and accumulate quiescent cell populations (Gonzalez et al. 2023). Spy1-CDK2 complex actively targets p27 for degradation and mediates epigenetic downregulation of other CDK inhibitors critical for neural differentiation (McAndrew et al. 2007), (Lubanska et al. 2021) and Spy1 protein levels accumulate to inhibit differentiation during normal mammary development (Golipour et al. 2008). Endogenous Spy1 levels elevate following acute sciatic nerve injury, suggesting a potential role in CNS regeneration (Huang et al. 2009). We demonstrated previously that, Spy1 is transcriptionally activated by c-Myc (Golipour et al. 2008) and Spy1 protein is stabilized following p53 deletion/mutation (Fifield et al. 2019). Consequently, protein levels of Spy1 are elevated in several types of cancer (Fifield et al. 2019; Lubanska and Porter 2014) and Spy1 regulates symmetric division and expansion of aggressive populations of brain tumour initiating cells in human glioma (Lubanska et al. 2014).

Using a conditional mouse model, this is the first study to characterize the consequence of elevated levels of cell cycle regulator, Spy1, in Nestin-positive NSC populations in the mammalian brain. We show that the induction of Spy1 regulates neuronal differentiation *in vitro* and *in vivo.* In primary cultures of NSCs, Spy1 promotes symmetric segregation of cell fate determinants and increases their susceptibility to oncogenic transformation. We further provide data to support that select drivers of glioma including depletion of functional p53 rely on endogenous Spy1 to maintain tumour progression *in vivo.* This work contributes to better understanding of the cell cycle mechanisms capable of tipping the balance of decisions in NSCs capable of predisposing this population of cells toward supporting oncogenesis.

## RESULTS

### Generation of pTRE-Spy1 transgenic mouse and NTA-Spy1 mouse model

To address the role of Spy1 in NSC populations of the mammalian brain, we generated a transgenic mouse system, termed NTA-Spy1, which permits inducible, spatially homogenous expression of Spy1 throughout the course of development as originally described by the Chodosh group (Gunther et al. 2002). The Flag-Spy1 sequence was removed from a previously described vector – Flag-Spy1A-pLXSN and was cloned into a pTRE-Tight-Caspase3 vector backbone (Figure S1A). A restriction digest using XhoI was performed to remove Flag-Spy1 coding sequence which was used for microinjection (Figure S1B). PCR analysis depicted pTRE-Spy1 positive founder mice through the presence of a 536bp band (Figure S1C). The pTRE-Spy1 mice express the open reading frame of Spy1 downstream of a promoter containing a series of tetracycline operator sequences. Lastly, the positive founder mice of a B6CBAF1/J background were found to transmit the pTRE-Spy1 transgene to their progeny (Figure S1D). To create the NTA-Spy1 mouse model, pTRE-Spy1 transgenic mice were crossed with Nestin-rtTA mice (Figure 1A). This mouse model uniquely allows the control of Spy1 expression under the Nestin promoter in the presence of doxycycline (dox).

**Figure 1.**
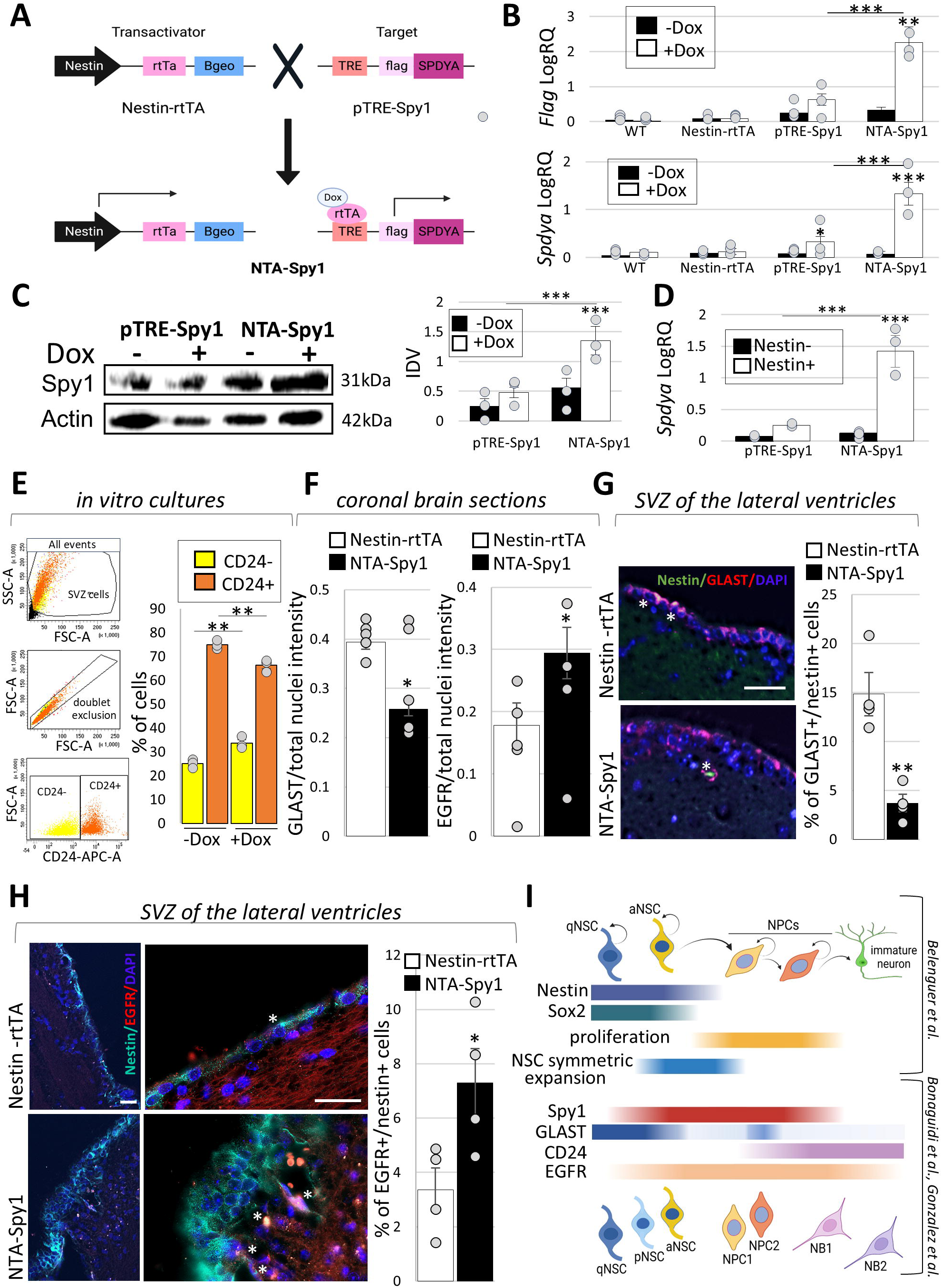
Characterization of NTA-Spy1 transgenic mouse (A) Schema presenting the generation of NTA-Spy1 mouse model. (B) QRT-PCR for Flag (top) and Spdya (bottom), WT: wild type (n=3, **p < 0.01, ***p < 0.001; Student’s *t*-test). (C) Western blot and densitometry analysis for Spy1 in neurosphere cultures. Integrated Density Values (IDV) Spy1/Actin (n=3, ***p < 0.001; Student’s *t*-test). (D) QRT-PCR for Spdya in sorted (FACS) Nestin-and Nestin+ cells (n=3, ***p < 0.001; Student’s *t*-test). (E) Flow cytometry analysis of CD24 in NTA-Spy1 cells (left) in dox treated (+Dox) and non-treated (-Dox) cultures, graphed as % of population tested (right). (n=3, **p < 0.01; Student’s *t*-test). (F) Quantified immunofluorescence of GLAST (left) and EGFR (right) in coronal brain sections Stain intensity to nuclei ratio graphed. (n=5, *p < 0.05; Student’s *t*-test). (G&H) Immunofluorescent staining to determine the number of GLAST+/Nestin+ (G) and EGFR+/Nestin+ cells (H) (*). Scale bar: 25 µm. Number of antibody+/Nestin+ cells graphed as ratio to nuclei number. (n=4, *p < 0.05, **p < 0.01; Student’s *t*-test). (I) A schema indicating cell type characteristics and specific marker expression during adult neurogenesis (Belenguer et al. 2021; Bonaguidi et al., 2011; Gonzalez et al., 2023). Created in https://BioRender.com. Results presented as mean values ±SEM. n represents separate replicates; number of individual mice tested per genotype.

### Characterization of the NTA-Spy1 mouse model

To evaluate the expression levels of Spy1 in the NTA-Spy1 model, mice were induced with dox staring *in utero*. NTA-Spy1 or control brains were micro-dissected to remove the SVZ at post-natal day 2 (PN2), and the cells were obtained through enzymatic tissue dissociation. To determine if Spy1 protein levels remain sensitive to alterations in dox, cells were cultured in the presence or absence of dox (referred to as ‘induction *in vitro* with dox’) and subjected to qRT-PCR and western blot analysis. NTA-Spy1-derived neurosphere cultures show significant increase in Flag and Spy1 expression when induced in *in vitro* with dox as compared to cells derived from all littermate controls (Figure 1B). NTA-Spy1 and pTRE-Spy1 littermate control cells treated *in vitro* with dox for 72 hours showed a 2-fold increase in Spy1 protein when compared to cells not treated with dox (Figure 1C). This permits the study of an increase in Spy1 protein levels within a single germline for a subset of assays. Using anti-Nestin antibody, cells were separated via fluorescence activated cell sorting (FACS) to determine if Spy1 levels were being upregulated specifically in the Nestin positive vs. negative populations using qRT-PCR.

Spy1 levels were increased in the Nestin positive populations within each genotype (Figure S1E&1D).

Previously published data using Nestin promoter-driven labeling suggested that the neural lineage origin is located in the Nestin+, Sox2+ NSCs, which undergo low frequency of symmetric division to maintain their pools and divide asymmetrically to generate fast proliferating neural progenitor cells, NPCs ^19^ (Fig. 1I). Recent cclassification based on the expression levels of confirmed astrocytic markers, such as glutamate-aspartate transporter (GLAST), along the assessment of the expression of the neuroblast marker, CD24, and the epidermal growth factor receptor (EGFR), allows for isolation of CD24-/GLAST^high^/EGFR^low/-^ quiescent NSCs (qNSCs) and CD24-/GLAST^low^/EGFR^+^ activated NSCs as well as subsets of NPCs; NPC1 (CD24^low/-^/GLAST-/EGFR+) and NPC2 (CD24^high^/GLAST+/EGFR+) and neuroblasts (NBs) (Fig.1I) (Gonzalez et al. 2023; Belenguer, Duart-Abadia, Domingo-Muelas, et al. 2021;Belenguer, Duart-Abadia, Jordán-Pla, et al. 2021). A series of flowcytometry experiments were conducted using cells derived from SVZs, at PN2, of uninduced NTA-Spy1 mice and maintained as neurospheres for at least 3 passages without addition of dox. Labelling with antibodies against Nestin, Sox2, CD24, GLAST and EGFR showed that the derived cultures contain over 80% of Nestin+ cells (Fig.S1F) and all Nestin+ cells express Sox2 in these conditions (Fig. S1G). Furthermore, the neurosphere cultures contain both CD24-and CD24+ cells (Fig.1E) with 94% of CD24-cells being GLAST negative and almost 98% CD24+ being GLAST positive (Fig.S1H). As more in-depth future analysis is warranted to confirm the identity of the isolated cultures, these results suggest that the primary neurospheres derived and maintained under the culture conditions in this study share a marker expression profile which is similar to that of activated NSCs (aNSCs) and/or neural progenitor cells (NPC1 and NPC2) (Belenguer, Duart-Abadia, Domingo-Muelas, et al. 2021; Belenguer, Duart-Abadia, Jordán-Pla, et al. 2021) (Fig.1I). NTA-Spy1 neurospheres were then treated with dox over three passages. This *in vitro* induction of Spy1 significantly upregulated % of CD24-cells while downregulating numbers of CD24+ cells when compared to cultures without dox (Fig.1E). Furthermore, in comparison to dox-untreated cultures, NTA-Spy1 dox-induced neurospheres showed a decrease of the percentage of GLAST+ cells and no significant difference of EGFR+ cells (Fig.1SI). Consistently, significantly fewer GLAST+ cells (Fig.1F; left) and Nestin+ cells enriched for GLAST expression (Fig.1G) were found in the adult NTA brain sections in mice of 5 months of age, exposed to dox starting *in utero* when compared to littermate controls. Analysis of the same sections revealed significantly higher overall levels of EGFR expression (Fig.1F; right) and upregulated number of EGFR-enriched Nestin+ cells (Fig.1H) in NTA-Spy1 versus control rains. These results suggest that upregulation of Spy1, in Nestin+ cells of the SVZ, potentially leads to a further increase of the pools of the activated NSCs.

### Enhanced proliferation and neurosphere formation in NTA-Spy1 primary cultures *in vitro*

Transient overexpression of Spy1 in wildtype neurospheres derived from the SVZ of neonatal mice reveals an increase in gene expression levels of key stem cell markers Musashi 1 (Msi1), CD133, Bmi1, Vimentin and Nestin as compared to empty vector control (Figure S2A). To assess the functional consequence of elevated Spy1 levels in the NSC population, mice were induced with dox starting *in utero* and SVZs were dissected from NTA-Spy1 mouse brains along with their littermate controls, between PN2 and PN4, and subjected to dissociation. The obtained cells were maintained and used in assays *in vitro* in the presence of 1μg/ml dox to assess the role of Spy1 in proliferation and self-renewal of the mouse-derived cultures. BrdU incorporation assay revealed that, in comparison to all controls, NTA-Spy1 cultures presented a significantly elevated percentage of BrdU+ cells over the total cell population scored (Figure 2A). Using a neurosphere formation assay, we found that NTA-Spy1 cells formed significantly more spheres over serial passages (Figure 2B), and that NTA-Spy1 neurospheres were significantly larger in size (Figure 2C&D) as compared to controls. To assess true clonal potential of NSCs derived from our model, single cell self-renewal assay was employed to eliminate potential aggregation factor in the assessment of neurosphere formation. Consistently with neurosphere formation assay, a higher proportion single-cell-derived spheres was quantified in NTA-Spy1 cultures in comparison to pTRE-Spy1 control (Figure 2E). qRT-PCR analysis of the neurospheres showed a significant increase in gene expression of stemness markers, CD133, Sox2 and Nestin, throughout serial passaging of NTA-Spy1 spheres compared to control (Figure 2F). NTA-Spy1 neurospheres showed also significantly elevated Sox2 protein levels when compared to pTRE-Spy1 control (Figure S3A). Further qRT-PCR analysis revealed that unlike pTRE-Spy1 littermate controls, NTA-Spy1 cells have upregulated expression levels of other reprogramming transcription factors, including Oct4, c-Myc, Klf4 and Nanog, in addition to Sox2 (Figure 2G). We then tested the gene expression levels of defined reprogramming transcription factors, Olig2, Sox2, Pouf3, Sall2, essential for propagating tumour initiating cells from differentiated progeny (Suvà et al. 2014) and found increased expression of all factors tested in cells derived from NTA-Spy1 mice in comparison to pTRE-Spy1 controls (Figure S3B). Due to an intrinsic increase in reprogramming genes, we wanted to determine whether NTA-Spy1 cells had the potential to sustain self-renewal regardless of being cultured in differentiation promoting media. We employed a re-sphering assay, as previously described (Kerosuo et al. 2008). Single cell suspension of 400 cells derived from neurospheres grown in EGF and FGF-containing were plated per well, in 96-well plates in the presence of dox and 2% FBS to promote differentiation (Figure S3C). At two weeks post differentiation, there was a four-fold increase of the percentage of cells forming new neurospheres, quantified, in the differentiation conditions in the NTA-Spy1 cohort as compared to pTRE-Spy1 control (Figure S3D). The percentage of NTA-Spy1 cells generating neurospheres in our assay was consistent with previously reported maximum re-sphering capacity of cells overexpressing Myc (1.5%, at two weeks) (Kerosuo et al. 2008). qRT-PCR analysis of differentiation markers S-100β (astrocyte marker), Foxo4 (oligodendrocyte marker), and Mapt (neuronal marker) showed an increase in their gene expression levels in the differentiated cultures compared to initial neurospheres followed by a decrease during re-sphering with significantly lower expression of all differentiation markers throughout all stages in NTA-Spy1 cultures relative to the pTRE-Spy1 control (Figure S3E). These data suggest that Spy1 regulates proliferation, self-renewal and differentiation in NSCs *in vitro*.

**Figure 2.**
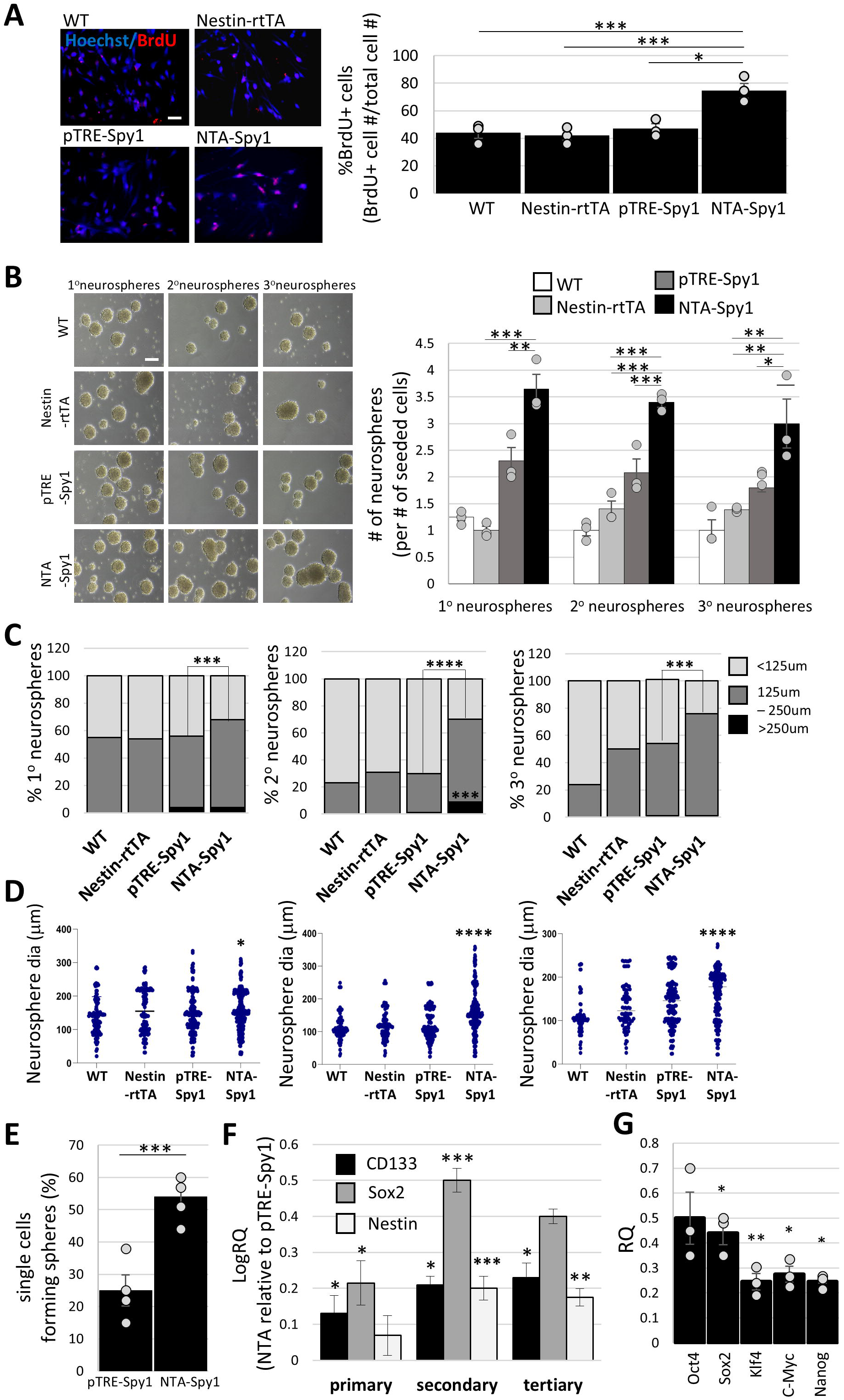
NTA-Spy1 derived cells have enhanced stemness characteristics (A) BrdU incorporation assay; representative images of cells (left). Scale bar: 50 µm. Data graphed as percentage of BrdU+ cells of total population, averaged over 5 fields of view/replicate (n=3, *p < 0.05, ***p < 0.001, one-way ANOVA). (B&C) Number (B) and diameter (C) of spheres scored over serial passages (primary-1°, secondary-2°, tertiary-3°) in a neurosphere formation assay. Scale bar: 100 µm. (n=3, *p < 0.05, **p < 0.01, ***p < 0.001, Student’s *t*-test (B), ***p < 0.0001; ****p < 0.00001, one-way ANOVA with Post Hoc Tukey HSD for pairwise comparisons between NTA-Spy1 and control (C). (D) Cumulative size distribution of serially passaged neurospheres (n=3, *p < 0.05, ****p < 0.00001, one-way ANOVA). (E) Clonal assay, percentage of single cells forming spheres quantified (n=4, ***p < 0.001; Student’s *t*-test). (F) QRT-PCR analysis for *CD133*, *Nesti*n and *Sox2* over serial passages of neurospheres. (n=4. *p < 0.05, **p < 0.01, ***p < 0.001, Student’s *t*-test). (G) QRT-PCR for *Oct4*, *Sox2*, *Klf4*, *c-Myc* and *Nanog*. RQ of NTA-Spy1 relative to control (n=3, *p < 0.05, **p < 0.01; Student’s *t*-test). Results presented as mean values ±SEM. n represents separate replicates; number of individual mice tested per genotype.

### NTA-Spy1-derived cells exhibit decreased neural differentiation capacity

To further determine the effect of increased levels of Spy1 on lineage differentiation, cells derived from SVZs of mice between PN2 and PN4 which had been continuously exposed to dox, starting *in utero*, were cultured in adherent conditions with continued exposure to 1μg/ml dox and 2% FBS to induce differentiation. Cell morphology was monitored over time (Figure 3A; left) and cells that contained neurite processes of at least two cell-body diameters in length were scored as differentiated, as described previously (Chuma et al. 2024), (Dravid et al. 2021). We found that NTA-Spy1 cells had a significant decrease in the percentage of differentiated cells compared to cells derived from control littermates (Figure 3A; right). To study the impact on specific neural differentiation lineages the cells were directed down astrocyte, oligodendrocyte, or neuronal lineages using validated differentiation protocols. Gene expression analysis demonstrated a significant decrease of S-100β and Foxo4 levels in NTA-Spy1 cultures compared to pTRE-Spy1 controls at 96 hours post differentiation (Figure 3B) and significant downregulation of the levels of both neuronal markers, beta-III Tubulin and Mapt, at earlier, 0-and 48-hour, timepoints (Figure 3B). Consequently, immunocytochemistry of beta-III Tubulin at 96 hours post differentiation of NTA-Spy1 cells in the presence and absence of dox (Figure S3F, left) revealed significantly lower levels of beta-III Tubulin in the induced versus un-induced cells (Figure S3F, right). We then analyzed the expression levels of p27 protein which plays a multifaceted role in neural differentiation (Domingo-Muelas et al. 2023; Nguyen et al. 2006) and is a target of Spy1-CDK2 complex (McAndrew et al. 2007). Using western blot analysis, we found that the levels of p27 protein were significantly lower in NTA-Spy1 compared to pTRE-Spy1 control cultures (Figure 3C). These experiments demonstrate that Spy1-mediated effects occur early during neuronal lineage differentiation time course and are concomitant with reduction in p27 levels.

**Figure 3.**
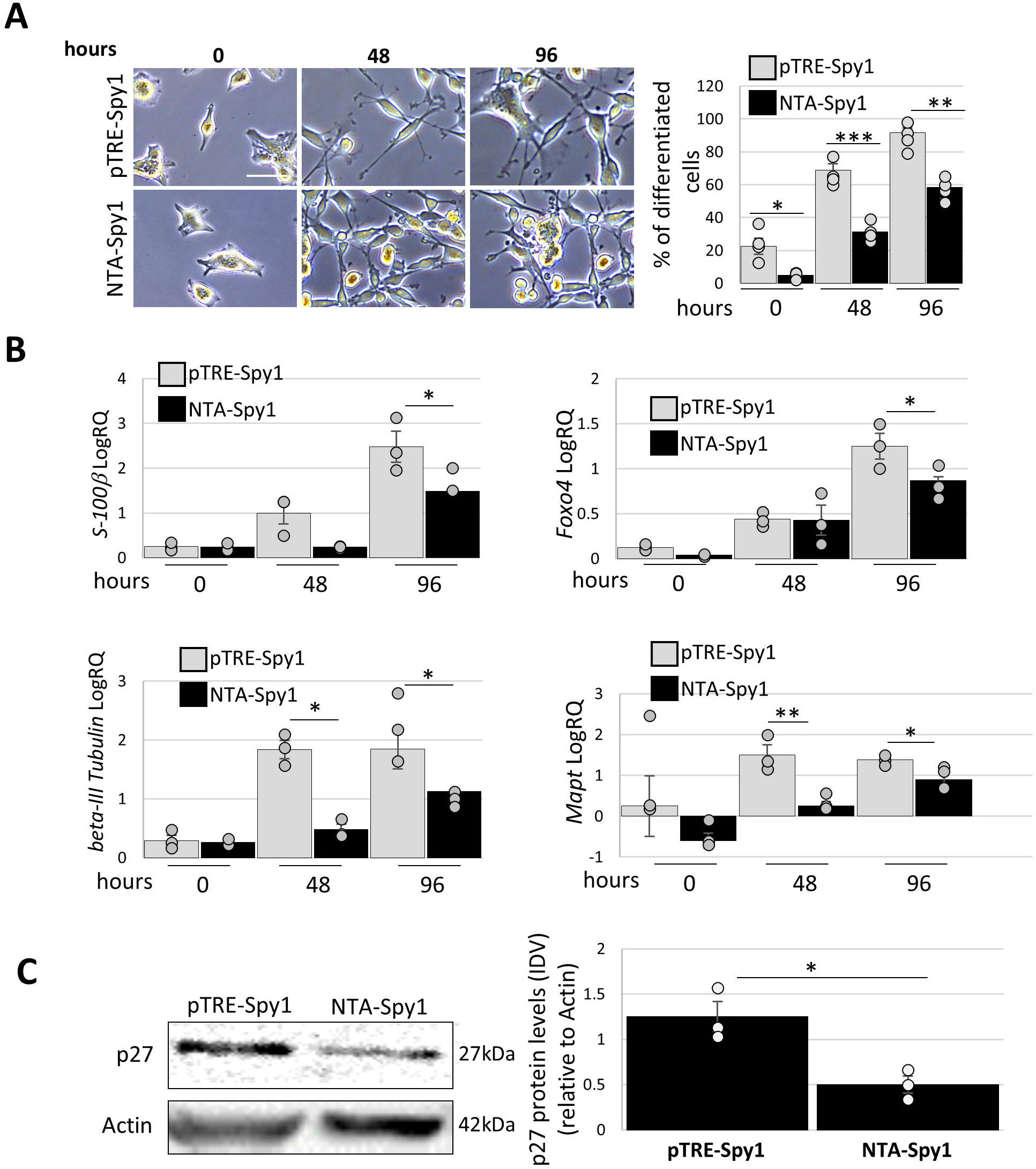
NTA-Spy1 cells decrease differentiation capacity (A) Differentiation time course in 2% FBS (left). Differentiated cells scored as percentage of total number of cells in the population analyzed (right). Scale bar: 30 µm. (n=4, **p < 0.01, ***p < 0.001; Student’s *t*-test). (B) QRT-PCR analysis for of *S-100*β, *Foxo4*, *beta-III Tubulin*, *Mapt* at the indicated time points of differentiation. (n=3, *p < 0.05, **p < 0.01; Student’s *t*-test). (C) Western blot (left) and densitometry analysis for p27. Protein levels scored as Integrated Density Values (IDV) and corrected by Actin. (n=3, *p < 0.05, **p < 0.01, ***p < 0.001; Student’s *t*-test). Results presented as mean values ±SEM. n represents separate replicates; number of individual mice/cultures tested per genotype.

### Regulation of the NTA-Spy1 cell fate depends on Musashi 1

We have previously demonstrated that Spy1 drives expansion of human glioma stem cells through regulating their mode of division (Lubanska et al. 2014). Asymmetric cell division supports the terminal differentiation identity of neurons in *C. elegans* (Bertrand and Hobert 2009). Although more evidence is still required to support existing data, it’s been suggested that similarly to *Drosophila,* in mice, the basic molecular mechanism of asymmetric cell division involves the subcellular localization of cell fate determinant protein, Numb (Shen et al. 2002). To study cell fate decisions of the NTA-Spy1 cells in comparison to littermate-derived controls, SVZs of mice at PN2, were first cultured in the presence of signals from the extracellular matrix, using Matrigel in the presence of dox (Figure S4A, left). Induction of Spy1 levels resulted in over 3-fold increase in the number of 3D colonies and a decrease in the percentage of differentiated cells (Figure S4A, right). Too determine potential impact of Spy1 on the distribution of Numb protein during the division of tested NSCs, the SVZ-derived cells were subjected to a cell pair assay, in the presence and absence of dox, followed by an immunofluorescent analysis of Numb levels between mitotic cell pairs. We found a significant decrease in asymmetric distribution of Numb protein in NTA-Spy1 cultures as compared to control littermate pTRE-Spy1 control (Figure S4B) which was consistent with changes to the distribution of CD133 between the tested cultures (Figure S4C).

The Notch pathway plays a vital role in NSC maintenance and self-renewal (Hitoshi et al. 2002) and it is regulated by Numb and Musashi l (Msi1) (Okano et al. 2002) (Figure S4D). Msi1 activation and function is directly induced through phosphorylation by Spy1-bound CDK2 (Arumugam et al. 2012). To investigate signaling effects of Msi1 in the NTA-Spy1 model, SVZ-derived cells were maintained as neurosphere cultures and assayed, in the presence of dox. Western blot analysis revealed that Msi1 protein levels are increased in NTA-Spy1 neurospheres as compared to pTRE-Spy1 (Figure S4E) which corresponded to significantly decreased levels of Numb protein (Figure S4E). To test whether Msi1 plays a role in Spy1-mediated regulation of Numb distribution, we used short interference RNA (siRNA) against Msi1. qRT-PCR analysis confirmed a significant decrease in Msi1 mRNA levels and a consequent decrease in Notch transcriptional targets, *Hes1* and *Hes5,* downstream of Msi1 (Figure S4F). The cells were then subjected to a cell pair assay, and we found that although there was a significant increase of the percentage of cell-pairs with symmetric distribution of Numb immunofluorescence signal in siControl NTA-Spy1 cultures compared to control, the mitotic call pairs with even Numb distribution were significantly downregulated in NTA-Spy1 siMsi1 treated cultures in comparison to siControl treatment (Figure S4G). Consequently, a cell pair assay with an antibody against activated Notch1 demonstrated significant increase in symmetric distribution of Notch Intra-Cellular Domain and a decrease of its asymmetric localization in NTA-Spy1 mitotic cell pairs treated with dox when compared to untreated control (Figure S4H). The obtained results show that overexpression of Spy1 increases symmetric distribution of cell fate determinant proteins, including Numb, and that this regulation is mediated by Msi1-Notch1 signaling. These data suggest a potential mechanism through which Spy1 could regulate the cell fate in NSCs.

### Long-term induction of Spy1 results in retained pools of NSCs and impaired cognitive functions

To assess the capacity of NTA-Spy1 mice to maintain long-term pools of NSCs, we tested, *in vitro* and *in vivo*, cell populations of the SVZs derived from 20-month-old NTA-Spy1 mice which had been continuously treated with dox, staring *in utero* and then maintained in 1μg/ml of dox as neurospheres. In comparison to littermate controls, NTA-Spy1 cells grown as neurospheres showed not only significantly increased proliferation, as assessed by BrdU incorporation assay (Figure 4A), but continued to demonstrate a block in differentiation at each time point, of the differentiation time course, tested (Figure 4B). Immunostaining of the obtained brain sections revealed an overall upregulation of the number of PCNA+ cells (Figure 4C) as well an increase in number of PCNA+ cells which were also Nestin positive (Figure 4D) in the region of SVZ in the brains of the NTA-Spy1 mice in comparison to control mice. *In vitro*, NTA-Spy1 cells showed increased mRNA levels of the proliferation marker Ki67 (Figure 4E), as well as decreased CKI gene expression levels of p21 (*Cdkn1a*), INK4 (*Cdkn2a*), and p53 (*Trp53*) (Figure 4F) as compared to pTRE-Spy1 cells. Histone γH2A.X is the most common marker for DNA double strand breaks and telomere shortening, and at times can also be used as a cellular senescence marker. Notably, immunofluorescence assay for γH2A.X expression showed that NTA-Spy1 cells had a significant decrease in the average number of γH2A.X foci per cell over five fields of vies compared to control littermates (Figure 4G). We also found a significant decrease in apoptosis as measured by Caspase 3/7 activity in the NTA-Spy1 cells compared to pTRE-Spy1 controls (Figure 4H). This suggests that long-term induction of the overexpression of Spy1 in NTA-Spy1 mice results in continuous proliferative activity in the neurogenic SVZ niche with no signs of NSC pool depletion.

**Figure 4.**
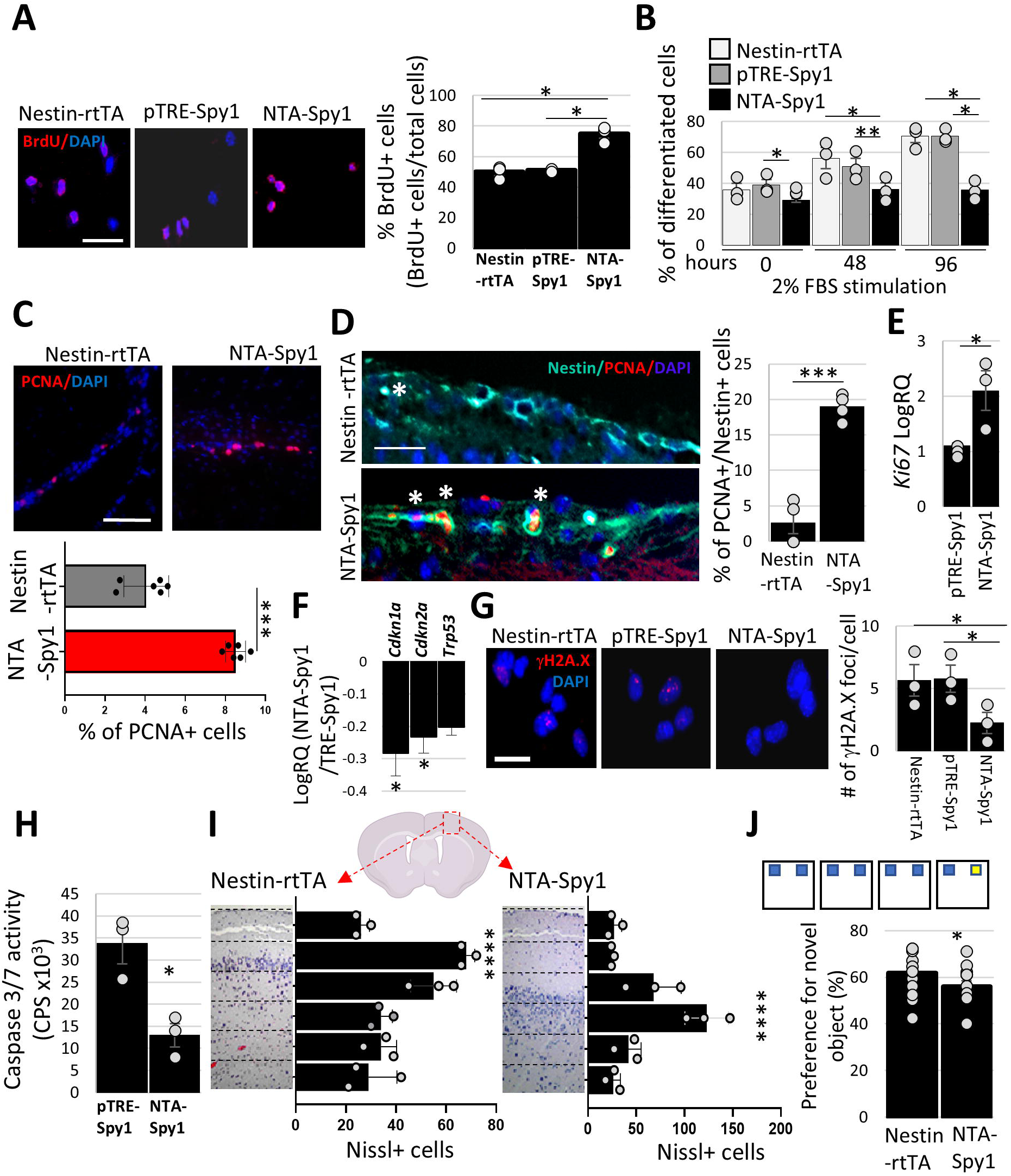
NSC pool is retained upon long-term induction of Spy1 in the NTA-Spy1 mouse brain (A) BrdU incorporation assay; representative images of cells (left). Scale bar: 50 µm. Data graphed as percentage of BrdU+ cells in population tested, averaged over 3 fields of view/replicate (n=3, *p < 0.05, Student’s *t*-test). (B) Differentiation time course in 2% FBS. Differentiated cells scored as percentage of total population analyzed, averaged over 3 fields of view/replicate (n=3, **p < 0.01, ***p < 0.001; Student’s *t*-test). (C&D) PCNA immunostaining in SVZ. Representative images (top). Scale bar: 100 µm. PCNA+ cells scored as % of cells in the field of view (bottom). (n=6, ***p<0.0001, two-way ANOVA). (D) PCNA/Nestin immunostaining in SVZ. Representative images (left). Scale bar: 20 µm. Nestin+/PCNA+ cells scored as % of cells in the field of view (right). (n=4, ***p < 0.001. Student’s *t*-test). (E&F) QRT-PCR analysis for *Ki67* (E) and *p21, INK4* and *p53* (F) in neurospheres (n=3, *p < 0.05, Student’s t-test). (G) γH2A.X immunostaining in cells. Representative images (left). Scale bar = 20 µm (n=3, *p < 0.05, Student’s *t*-test). (H) Caspase 3/7 luminescent activity assay in neurospheres (n=3, *p < 0.05, Student’s *t*-test). (I) Nissl stain analysis in coronal sections of the brain. Nissl+ cells were scored/region (dotted lines). (n=3, ****p<0.0001, one-way ANOVA). (J) Novel object recognition memory test. Schematic (top) and time spent at novel object versus the overall time mice spent at objects on day 4 (bottom) (n=12, *p < 0.05; Student’s *t*-test). Results presented as mean values ±SEM. n represents separate replicates; number of individual mice/cultures/sections tested per genotype.

To determine whether the sustained self-renewal of NSCs in NTA-Spy1 mice relative to controls affects phenotypic characteristics of corticogenesis, we stained coronal brain sections of the 20-month-old Nestin-rtTA and NTA-Spy1 mice which had been continuously exposed to dox, starting *in utero*, with a neuron specific stain, Nissl, and scored Nissl+ cells in the selected regions of the cortex. NTA-Spy1 mice showed a shift in neuronal positioning of the cortical plate (Figure 4I) with obvious accumulation of neuronal bodies in closer proximity to ventricular zone as compared to Nestin-rtTA control mice. Consistently, an immunofluorescent staining using an antibody against Neuron Specific Nuclear Protein (NeuN) revealed increased density of mature postmitotic neurons in layer IV of the cortex in brain sections obtained from NTA-Spy1 mice in comparison to littermate controls (Fig.S5A). Using additional brain sections, we then investigated the expression of Pou3f2 and neurofilament H (NFH), proteins specific to neurons of the upper and deep cortical layers, respectively (Fig. S5B&S5C). In comparison to pTRE-Spy1 control, NTA-Spy1 mice demonstrated significantly elevated accumulation of neurons positive for Pou3f2 in cortical layer V (Fig. S5B). The expression levels of NFH were quantified as fluorescent signal intensity which was significantly downregulated across all the cortical layers of the NTA-Spy1 in comparison to control brain sections with the most significant decrease of NFH in the cortical layer VI (Fig. S5C). These results show that the overexpression of Spy1 throughout the brain development leads to changes to the density and distribution of upper and deeper neuronal types suggesting role of Spy1 in the regulation of corticogenesis in the mammalian brain.

To test whether the observed changes to neuronal positioning were affecting cognitive functions in the tested mice, we performed the novel object recognition task, as demonstrated by the schematic (Figure 4J; top). NTA-Spy1 mice had significantly decreased preference for novel objects brought to the task set up as compared to their littermate controls (Figure 4J; bottom). In summary, upregulation of Spy1 levels, in Nestin+ cells, results in changes to density and composition of the cortical layers and impairs cognitive functions in NTA-Spy1 mice.

### Spy1 alters expression of genes associated with neural malignancies in NTA-Spy1 model

Given the established role of Spy1 in several types of cancer including malignancies of neural origin, we investigated whether the continuous induction/overexpression of Spy1 in the NTA-Spy1 mice could play a role in transformation of NSCs in face of applied pro-oncogenic changes. We first studied the effects of the increased levels of Spy1 on the expression pattern of selected tumour suppressor genes, p53 and Pten, in the brains of 4-month-old mice on continuous dox diet beginning *at utero*. Immunostaining of the brain sections with anti-p53 antibody showed a small, yet statistically significant increase in the number of cells staining positively for p53 in the hippocampus of NTA-Spy1 mice compared to pTRE-Spy1 mice, while no difference was observed in cells of the SVZ or cortex (Figure S6A&S6C). A statistically significant increase in the percentage of cells staining positively for Pten expression was scored in the hippocampus and cortex of NTA-Spy1 mice compared to control, with no difference observed in the SVZ (Figure S6B&S6D). These findings suggest that tumour suppressor genes may increase to counteract the effects of elevated Spy1 in subsets of cells. To further explore this, we downregulated the levels of p53 via shRNA and inhibited Pten using bpV(HOpic) inhibitor (PTENi) in primary, 1μg/ml dox-induced neurosphere cultures from the 4-month-old NTA-Spy1 and control pTRE-Spy1 mice (Figure S6E&F). Gene expression analysis revealed that there was an overall significant increase in Cyclin D1 and c-Myc levels in the NTA-Spy1 in comparison to the pTRE-Spy1 cultures (Figure S7A). The expression levels of selected targets were further elevated with downregulation of p53 and inhibition of Pten individually and in combination in NTA-Spy1 derived cells compared to the pTRE-Spy1 control (Figure S7A). This data supports that overexpression of Spy1 protein levels correlate with alterations in other genes/proteins associated with increasing proliferation and stemness found in brain malignancies.

### Spy1 enhances oncogenic transformation in neurosphere cultures in the presence of selected aberrant events

A hallmark characteristic of oncogenic transformation is the ability to survive and proliferate in low attachment conditions (Figure S7B) (Rotem et al. 2015). Cultures derived from dissected SVZs of NTA-Spy1 or pTRE-Spy1 mice at 4 months of age, which have been dox-induced starting *in utero*, were transduced with shRNA against p53 and/or RasV12, using lentivirus, and treated with PTENi or vehicle control. The cells were then cultured in the presence and/or absence of dox in low adhesion conditions using ultra-low attachment plates for five days and their viability was assessed using 3D Cell Titer Glo assay. We found that pTRE-Spy1 littermate controls, regardless of the presence of dox, had significantly higher viability when grown in attachment conditions which is consistent with the growth preference of non-transformed cells (Figure S7C, top). However, dox-induced NTA-Spy1 cells exhibited enhanced viability in both growth conditions (Figure S7C, bottom), suggesting enhanced survival capacity of NTA-Spy1 cultures in non-adherent conditions. The viability of NTA-Spy1 cultures was significantly enhanced in ultra-low attachment conditions in the presence of overexpression of RasV12 and/or combined with the treatment with PTENi and siRNA against p53 (Figure S7E), this was not seen with inhibition/knockdown of Pten or p53 alone (Figure S7D). We then conducted classical soft agar colony formation assay in NTA-Spy1 cells compared to pTRE-Spy1 controls in the presence of dox with or without combined attenuation of p53 and Pten. At 21 days post-seeding, colonies with a diameter equal to or greater than 50 microns were observed in all conditions tested except pTRE-Spy1 culture without tumour suppressor downregulation in which NTA-Spy1 successfully generated colonies (Figure 5A). The transformation potential of the cells, in face of p53 and Pten manipulation, was assessed by subtracting the number of colonies obtained in the condition with no tumour suppressor manipulation from the following treatments (Figure 5B). There was also a significant increase in the number of colonies in the NTA-Spy1 in comparison to pTRE-Spy1 upon p53 knockdown and PTENi treatment, alone. Significant increase in the number of colonies was observed in NTA-Spy1 NSCs which had both the shp53 and PTENi combined (Figure 5B). We then conducted a colony formation assay using NTA-Spy1 and pTRE-Spy1 cells transduced with RasV12 and/or control. Although the overexpression of Ras resulted in colonies forming regardless of Spy1 overexpression, the NTA-Spy1 NSCs generated significantly more colonies than pTRE-Spy1 cells (Figure 5C). It has previously been shown that Spy1 overrides the DNA damage response in a variety of cell types (Fifield et al. 2019; Gastwirt et al. 2006). To determine whether NTA-Spy1 cells derived from SVZ have altered response to DNA damage, cells were treated with a chemotherapy drug, temozolomide (TMZ), relied upon in the treatment of many neural tumours, along dox treatment. NTA-Spy1 cells demonstrated significantly lower levels of apoptosis as measured by Caspase 3/7 activity assay in comparison to Nestin-rtTA control cells (Figure 5D). The treatment resistance was further enhanced in NTA-Spy1 cells when p53 knockdown was introduced (Figure 5E). The response to treatment in NTA-Spy1 cells was then tested in the context of overexpression of an oncogene and we found that increased levels of RasV12, contributed to further resistance to cell cycle arrest mediated by a synthetic CKI, NU2058, and/or TMZ, alone and in combination in NTA-Spy1 cells compared to pTRE-Spy1 control (Figure 5F). To finally address the capacity of Spy1 to support tumour formation in face of the oncogenic changes we employed *in vivo* models. First, using zebrafish xenograft model, we were able to engraft small numbers of NTA-Spy1 cells or littermate pTRE-Spy1 controls, which were pre-injection treated with siRNA against p53 (si-p53) or si-control, into zebrafish larvae at 48 hours post fertilization (hpf). Dox was added to fish water 8 hours post injection and the number of resulting foci per larvae was scored 4 days after (4dpi) (Figure 5G, left). Si-p53 increased foci in all conditions over control and there was a significant increase in foci formation in si-p53 NTA-Spy1 cells over si-p53 pTRE-Spy1 control (Figure 5G, right). Syngeneic GL261 mouse glioma model which is characterized by driving mutations in p53 and Ras, was then employed to further investigate the importance of Spy1 in supporting brain tumourigenesis. GL261 cells were transduced with shRNA against Spy1 or a scrambled control. The cells were sorted via FACS to enrich for solely transduced cells which were then transplanted orthotopically into neonatal C57BL/6 mice of syngeneic background. 14 days post injection 7/8 mice injected with the control developed visible tumours whereas tumours from shSpy1 cells were found in 4/8 mice at the 14-day timepoint (Figure 5H, left). Analysis of the tumour burden at 15 days post injection showed significantly downregulated brain tumour size in mice injected with shSpy1 cells when compared to the scrambled control (Figure 5H, right). Given slower growth of the tumours, analysis of the tissue sections revealed increased levels of senescence-associated β-galactosidase in pLB-shSpy1 tumours compared to control tumours (Figure 5I). qRT-PCR analysis of genes indicative of senescence showed upregulation of the mRNA levels of p21 and p16 in pLB-shSpy1 compared to scrambled control tumours (Figure 5J). The data suggest that Spy1 not only cooperates with known aberrant changes found different types of neural cancers to enhance the oncogenic potential of normal NSCs but also contributes to proliferation/growth in tumours driven by p53 and Ras mutations.

**Figure 5.**
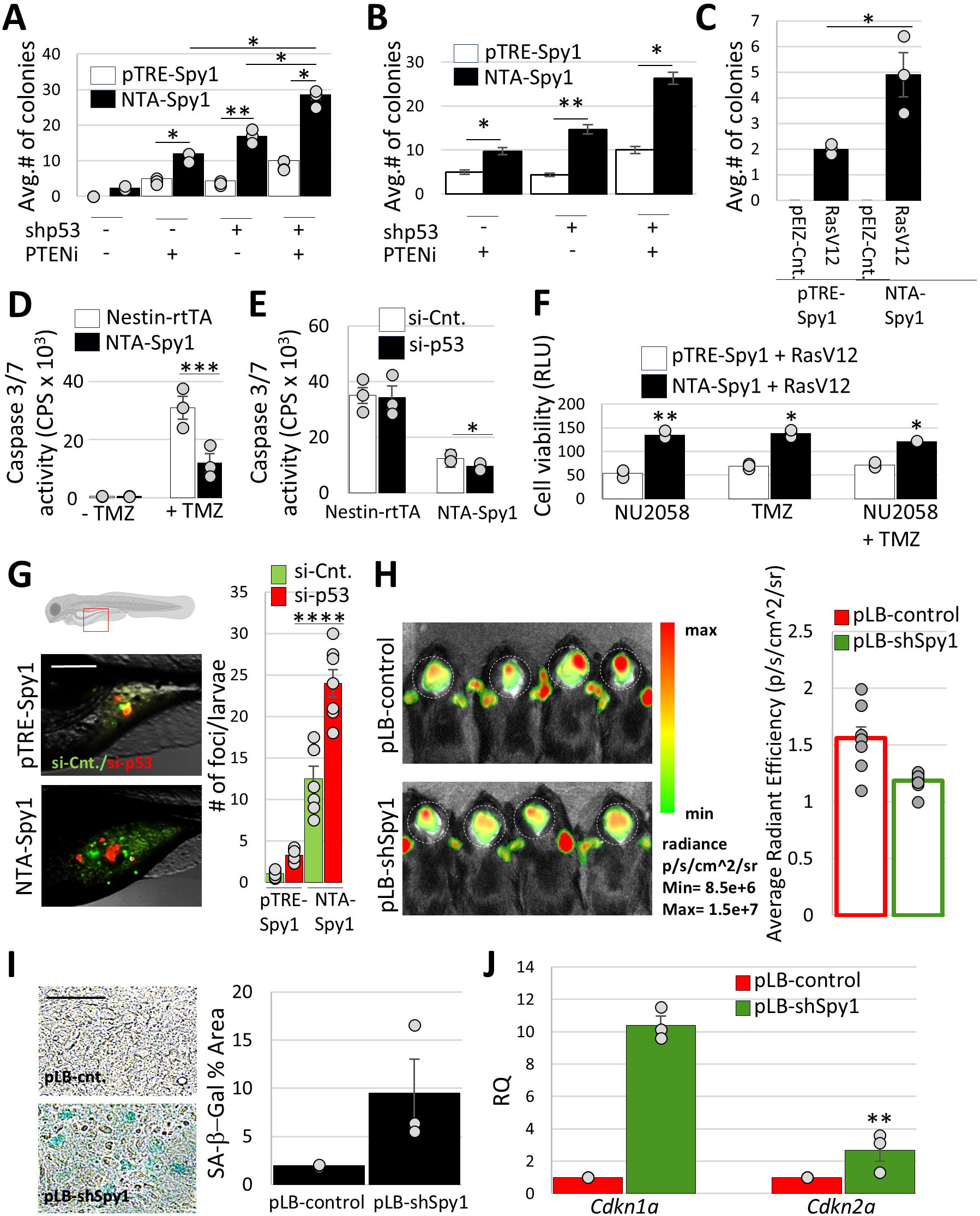
Spy1 plays a role in oncogenic transformation of NSCs (A&B) Soft agar colony formation assay. Colonies scored and graphed as average number of colonies per condition (A) and number of colonies in all conditions corrected by subtracting of the number of colonies obtained under control conditions-without p53 and/or Pten manipulation. (n=3, *p < 0.05, **p < 0.01, one-way ANOVA). (C) Soft agar colony formation assay in cultures transduced with pLENTI-CMV-RasV12/pEIZ-Control vector (n=3, *p < 0.05; Student’s *t*-test). (D&E) Caspase 3/7 luminescent activity assay in neurospheres treated with 25μM TMZ (+TMZ) and/or vehicle control (-TMZ) (D) and with siRNA control (si-Cnt.) or si-p53 in the presence of 25μM TMZ (E). (n=3, *p < 0.05, ***p < 0.001, Student’s *t*-test). (F) Luminescence-based 3D Cell Titer Glo Viability assay. Cells treated with 30μM NU2058 and/or 25μM TMZ alone or combined, Relative Luminescence Units (RLU). (n=3, *p < 0.05, **p < 0.01; Student’s *t*-test). (G) Zebrafish larvae xenograft assay. Representative images at 4dpi (left). Scale bar = 375 μm. siRNA control (si-Cnt). Foci > 50 μm in diameter scored per zebrafish larvae using ImageJ software (n=7, ****p < 0.0001, one-way ANOVA). (H) Tumour burden analysis. pLB-scrambled control (pLB-control) or pLB-shSpy1-transduced GL261 cells injected orthotopically. Fluorescence intensity quantified using Aura software as Average Radiant Efficiency (n=8, *p < 0.05, ***p < 0.001; Student’s *t*-test). (I) SA-β-Gal activity assay in pLB-control and pLB-shSpy1-derived tumours. Scale bar = 40 μm. SA-β-Gal activity graphed as percentage of stained area/section. (n=3, technical replicates from one tumour). (J) QRT-PCR analysis for of *Cdkn1a* and *Cdkn2a* in tumour tissue (n=3, technical replicates from one tumour).

## DISCUSSION

A study by Gonzalez *et al*. demonstrated the essentiality of the atypical cell cycle regulator Spy1 in the activation of NSC pools from the quiescent state followed by decreased neurogenesis (Gonzalez et al. 2023). We have demonstrated previously that Spy1 plays a critical role in symmetric division and clonal expansion of brain tumour initiating cells obtained from glioma patients (Lubanska et al. 2014) and that Spy1-CDK2 complex regulates reprogramming efficiency to a pluripotent state ^11^. The molecular mechanisms by which Spy1 regulates the NSC pool in the normal adult brain remain fundamentally unknown.

This study established the NTA-Spy1 mouse model which successfully permitted Tet-regulated, spatial control of the expression of Spy1 under the Nestin promoter allowing for detailed study of Spy1 mediated cell cycle effects throughout the brain development and in response to potential malignant changes in the pools of selected NSCs. Transactivation Tet-ON system, used in this study is characterized by basal leakiness resulting in increased-basal expression levels of the protein of interest. To control for this, where pTRE-Spy1 mice were used as controls we incorporate other littermate controls and the effects/differences observed in the NTA-Spy1 mice/cells which were significant in comparison to the wild type and Nestin controls were also significant when compared to pTRE-Spy1 validating proper control of our experiments. Several experiments in this study were conducted using neurosphere cultures of primary cells derived from our model. High enrichment in Nestin (∼80%) and Sox2 (∼100%) in our neurosphere cultures supports selection for NSCs by the methods utilized in this study (Bonaguidi et al. 2011). Although further characterization of NSCs revealed marker expression profile which correlated with that of activated NSCs, NPC1 and NPC2 (Belenguer, Duart-Abadia, Domingo-Muelas, et al. 2021; Belenguer, Duart-Abadia, Jordán-Pla, et al. 2021) and was consistent with previously demonstrated expression of Spy1 in these populations (Gonzalez et al. 2023), our study is not utilizing a subset of other markers including Lin+/-, CD45+/-, CD31+/-, Ter119+/-, CD9+/-required in the extensive depletion/enrichment protocol (Belenguer, Duart-Abadia, Domingo-Muelas, et al. 2021; Belenguer, Duart-Abadia, Jordán-Pla, et al. 2021) and thus cannot be used in direct comparison to NSC types discussed by others just yet.

Our *in vitro* data supports that Spy1 increases proliferation while maintaining stem cell attributes of the Nestin+ population in the brains of both young and aging mice. Increased levels of Spy1 in neurosphere cultures enhance stemness capacity and prevent functional differentiation down all lineages with enhanced impact on neuronal lineage. NTA-Spy1 mice demonstrate a subtle impairment in recognition memory which is coincident with an altered relative composition of the cortical plate layers with deeper layers accumulating markers of the upper cortex and all layers presenting dramatic downregulation of the deeper neuronal markers. Given the established “inside out” model of cortical development (Fan et al. 2008) our data suggest a potential role of Spy1 in the control of differentiation and pool size of early and late migrating neurons, respectively. Consistently, during corticogenesis, nestin positive progenitors demonstrate temporal specificity along developmental stages with early progenitors contributing to all neuronal layers while late progenitors establish upper cortical layers (Burns et al. 2009). These studies support our data and hypothesis that Spy1-mediated impact on neuronal density and composition of the mammalian cortex is timing dependent where Spy1 blocks terminal differentiation of early neurons forming deeper layers and expands the pools of progenitors which contribute late/upper layer neurons. More in depth investigation, however, is necessary to further dissect, characterize and validate this role for Spy1.

Our work proposes a role for Spy1 in the induction of a shift in NSC fate decisions which are regulated by symmetric over asymmetric distribution of cell fate determinants, including Numb. Although functional consequences of the accumulation of these proteins in the analyzed mitotic pairs require additional investigation, we speculate that potential shift towards symmetric mode of division can underlie the observed long-term self-renewal of NSCs, and potential expansion of progenitor pools involved in corticogenesis in NTA-Spy1 mice. We demonstrate that Spy1-mediated distribution of Numb depends on Msi1 signaling and Msi1-Numb-Notch axis has been also implicated previously in malignant transformation (Forouzanfar et al. 2020). Previously published studies showed that long-term/aging populations of NSCs accumulated dangerous changes in critical genes guarding their integrity and persistence of those changes results in tumourigenesis (Talma et al. 2021). Although, we show that combined downregulation of p53 or Pten tumour suppressor levels and/or overexpression of potent oncogenes, with consistently elevated levels of Spy1 in normal NSCs increases oncogenic transformation *in vitro*, a fine-tuned analysis of pro-oncogenic changes cooperating with Spy1 to drive tumourigenesis *in vivo* is warranted. We show that downregulation of Spy1 levels results in decreased tumour burden from orthotopic injection of glioma cells carrying abrogated p53 function and Ras protein mutation which is in agreement with our previous findings showing that degradation of Spy1 is an essential component of p53-mediated tumour suppression (Fifield et al. 2019). Of importance is the fact, that upon characterization of those tumours we find an increase in characteristics indicative of increased senescence in tumours with downregulated levels of Spy1 which suggest that Spy1-mediated effects in that system are instructive in nature.

In summary, this work demonstrates that upregulation of Spy1 under the Nestin promoter results in increased proliferation and expansion of NSCs characterized by symmetric accumulation of cell fate determinants in the mitotic pairs, in Msi1-dependent manner. These Spy1 mediated effects lead to decline in neurogenesis, altered corticogenesis and increased susceptibility to malignant transformation in face of selected pro-tumourigenic changes.

## EXPERIMENTAL PROCEDURES

### RESOURCE AVAILABILITY

#### Lead Contact

Further information and requests for resources and reagents should be directed to and will be fulfilled by the lead contact, Lisa Ann Porter.

#### Materials Availability

The mouse lines used in this study are available to academic groups upon signing of an MTA.

#### Data and Code Availability

- Any additional information required to reanalyze the data reported in this paper is available from the lead contact upon request.
- This study does not report original code.

#### Mouse dox diet and induction

Mice were fed with Rodent diet (2018, 2000 Dox, G) (Harlan, #TD.09633) containing 2018, Teklad Global 18% Protein Rodent Diet and 2g/kg of Dox Hyclate. Unless specified otherwise, all mice used in the experiments including primary cell extraction were induced at the time of conception (*in utero*) as females were fed with the diet ad libitum starting at the time of breeding.

#### Primary Cell Culture

Primary cells were cultured as neurospheres with addition of dox at 1μg/mL at the time of post-extraction seeding, in growth media consisting of serum-free Neurobasal™ medium (Thermo Fisher Scientific, #21-103-049) containing B-27™ supplement (Thermo Fisher Scientific, #17504044), human epidermal growth factor recombinant protein (hEGF) (20 ng/mL; Thermo Fisher Scientific, #PHG0311), human fibroblast growth factor-basic recombinant protein (hbFGF) (10 ng/mL; Sigma-Aldrich, #F0291), and 2 mM L-glutamine (Thermo Fisher Scientific, #25030081) at 37°C and 5 CO_2_ in ultra-low attachment plates (ULAPs). Neurospheres were subcultured every 5-10 days and culture media were replenished every 48-72 hours with 1μg/mL dox addition.

For all *in vitro* experiments, the induction continued by adding dox to a final concentration of 1μg/mL at the time of experiment set up and was readded with media change every 48-72 hours through the time of the experiment.

## ACKNOWLEDGEMENTS

We would like to thank staff in the animal care facility at the University of Windsor as well as Dr. Elizabeth Fidalgo da Silva for technical support. We acknowledge the funding from Faculty of Science, WE-SPARK Health Institute, and Office of the VPRI at the University of Windsor for support of the Flow Cytometry Core Facility. IQ, AR and KFS acknowledge support from the Queen Elizabeth II Ontario Graduate Scholarship. EB acknowledges support from the Natural Sciences and Engineering Research Council of Canada for Undergraduate Student Research Award. This work was supported by operating funds from the Canadian Institutes Health Research to LAP (Grant #406811).

## AUTHOR CONTRIBUTIONS

IQ, DL, FKS, HPM, SA, EB, EF, AR, MA, SD and DLi carried out the experiments presented in this manuscript. DL, IQ, FKS, BF and LAP supported experimental design. IQ, DL, BF, JB, HPM supported derivation and maintenance of the mouse colony. DL, IQ, AR, SK supported data analysis. IQ, DL, FKS, AC, HPM, SA, JB, BF, and LAP supported the writing of this manuscript. Grant funding to LAP supported this project. All authors edited the manuscript and provided comments on the intellectual content.

---

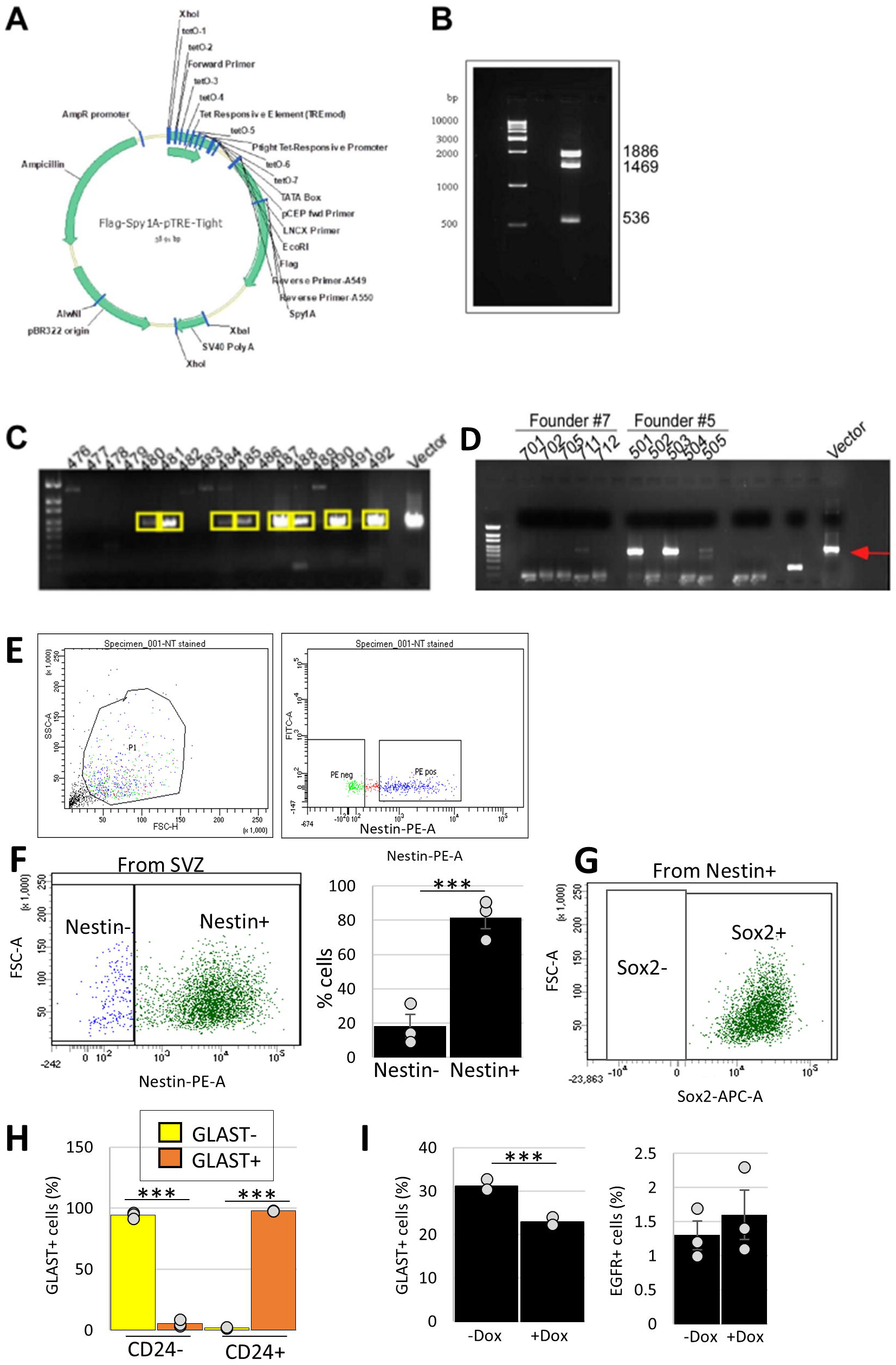

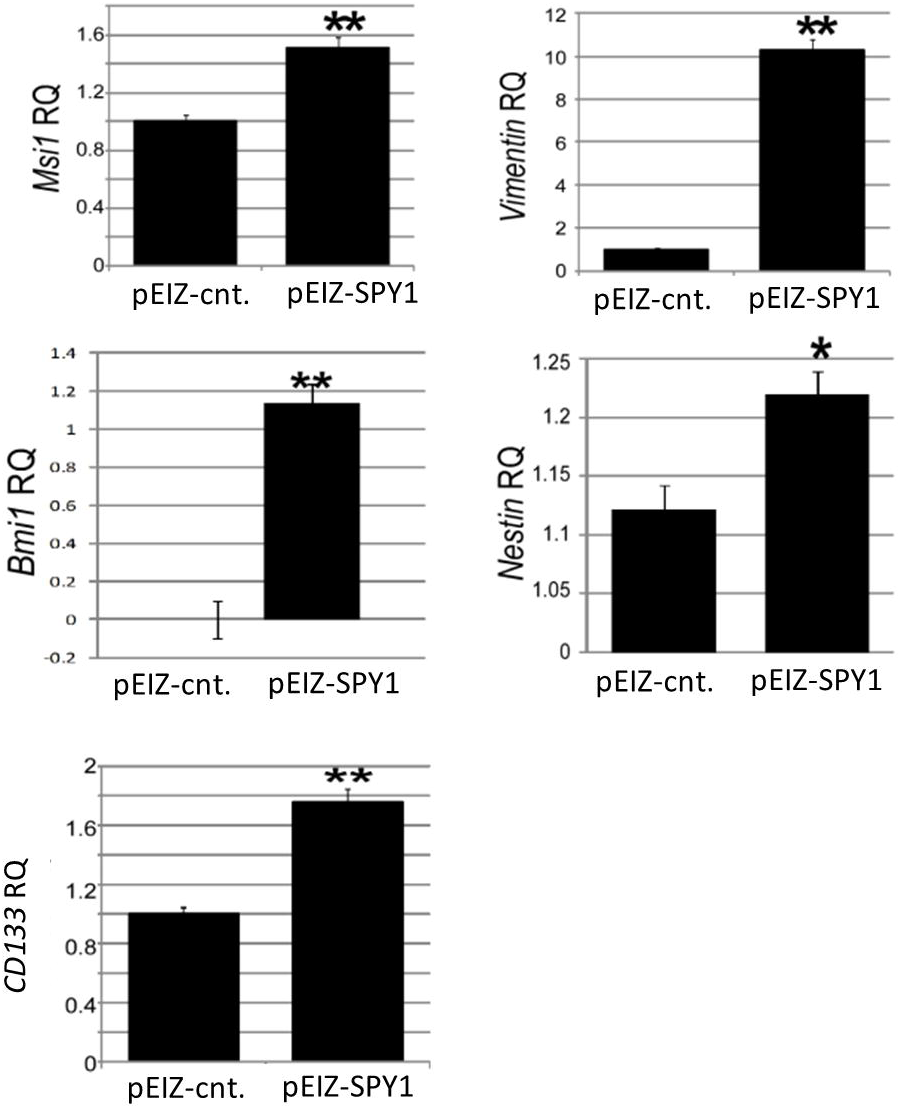

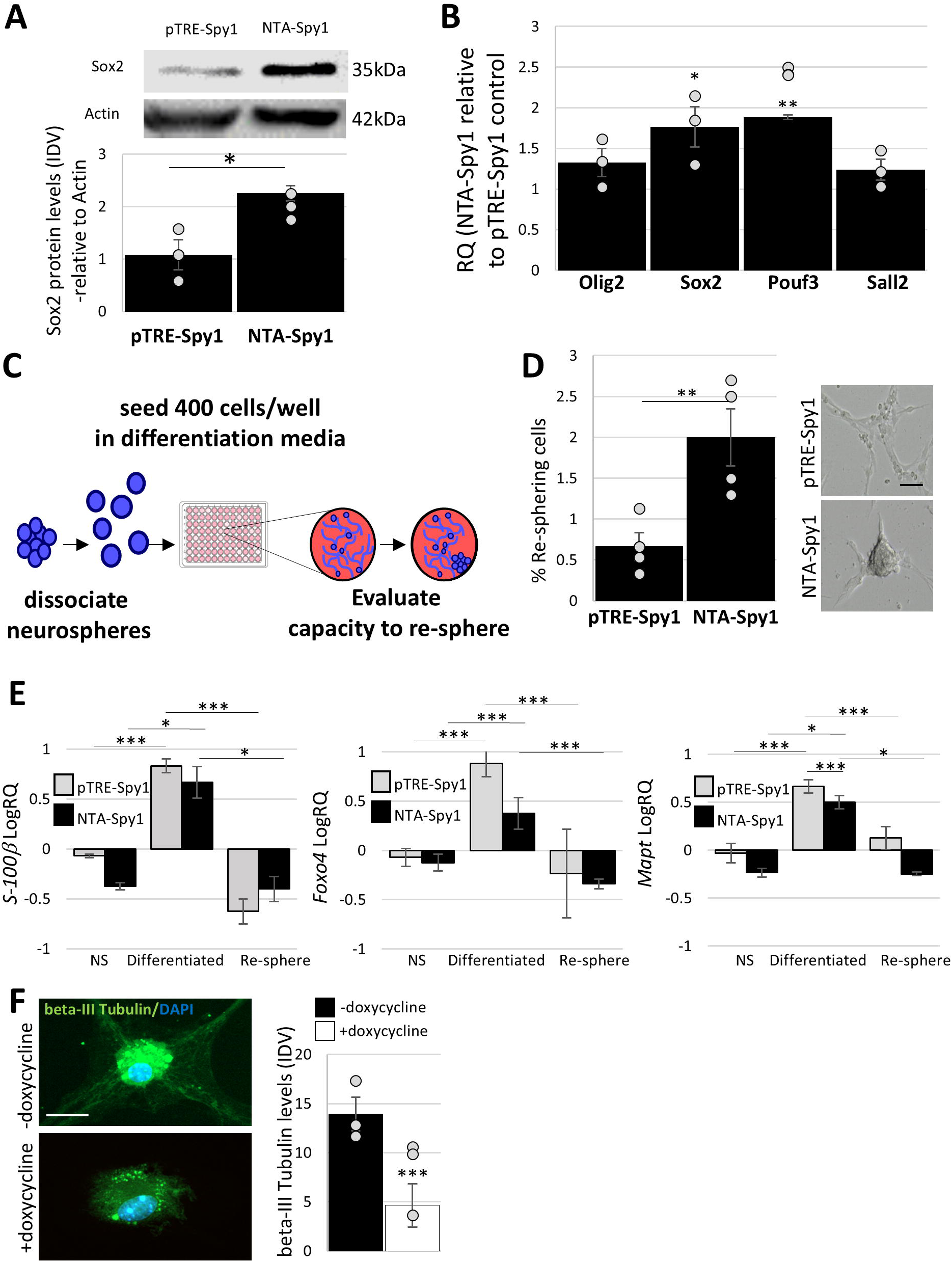

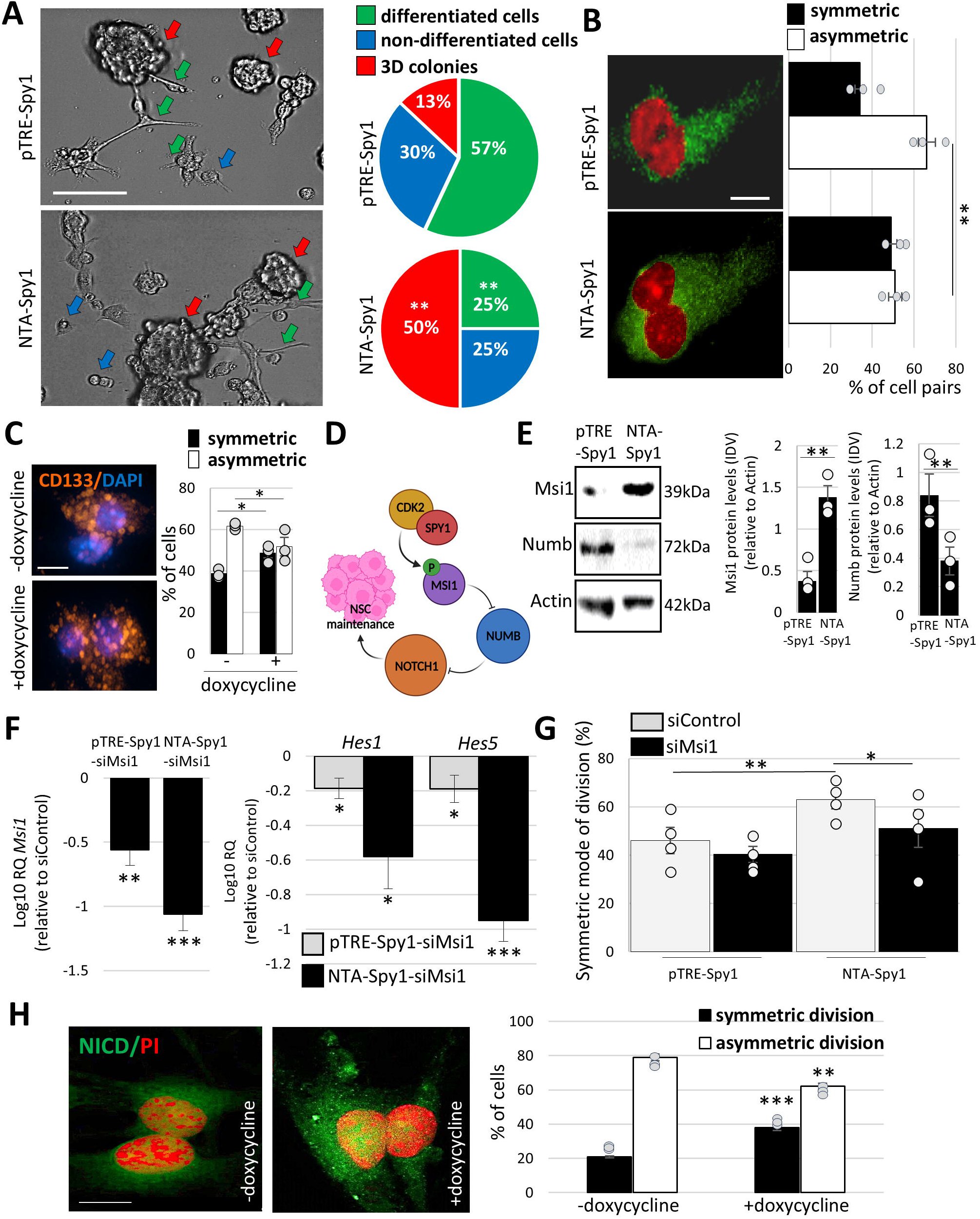

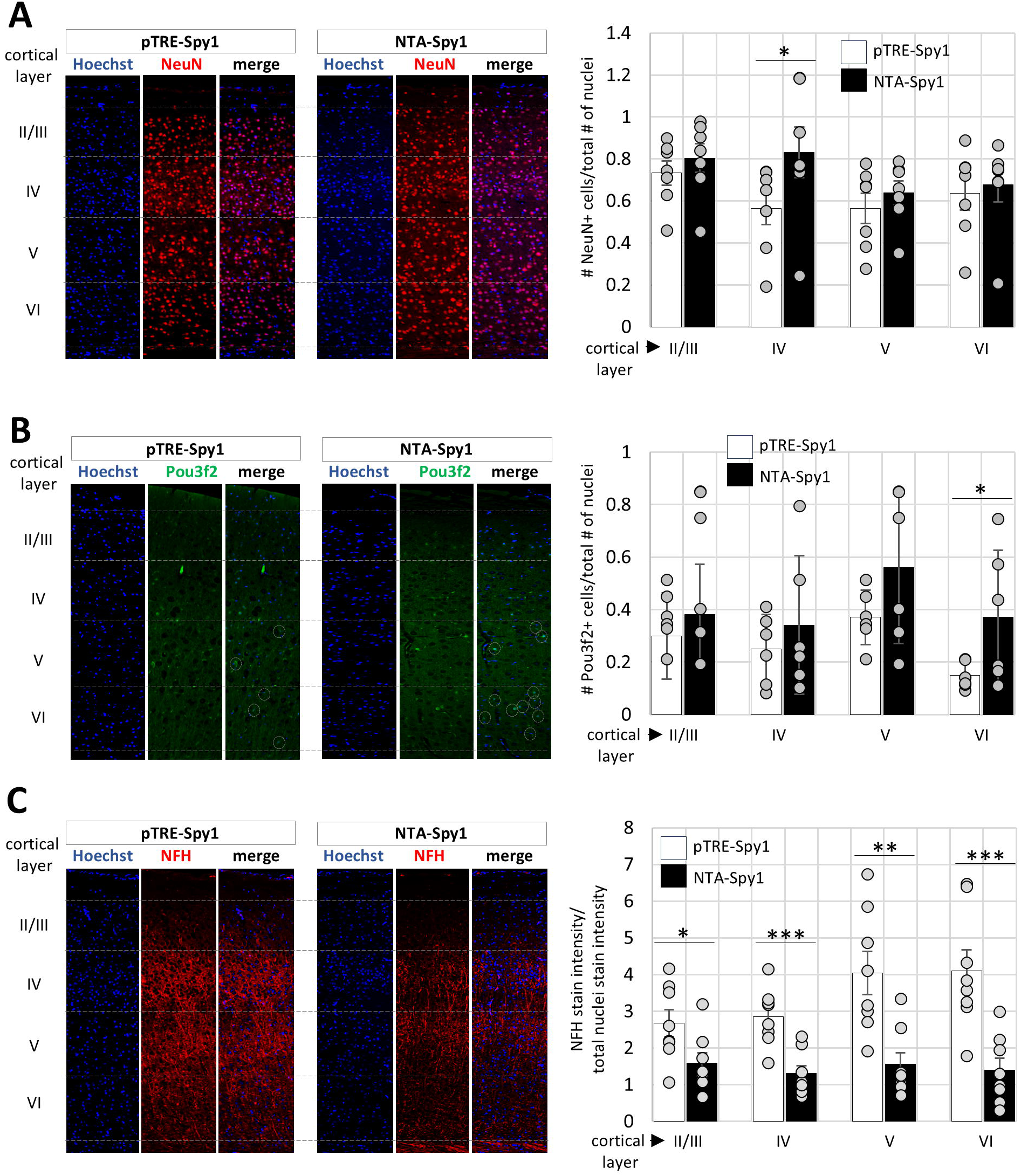

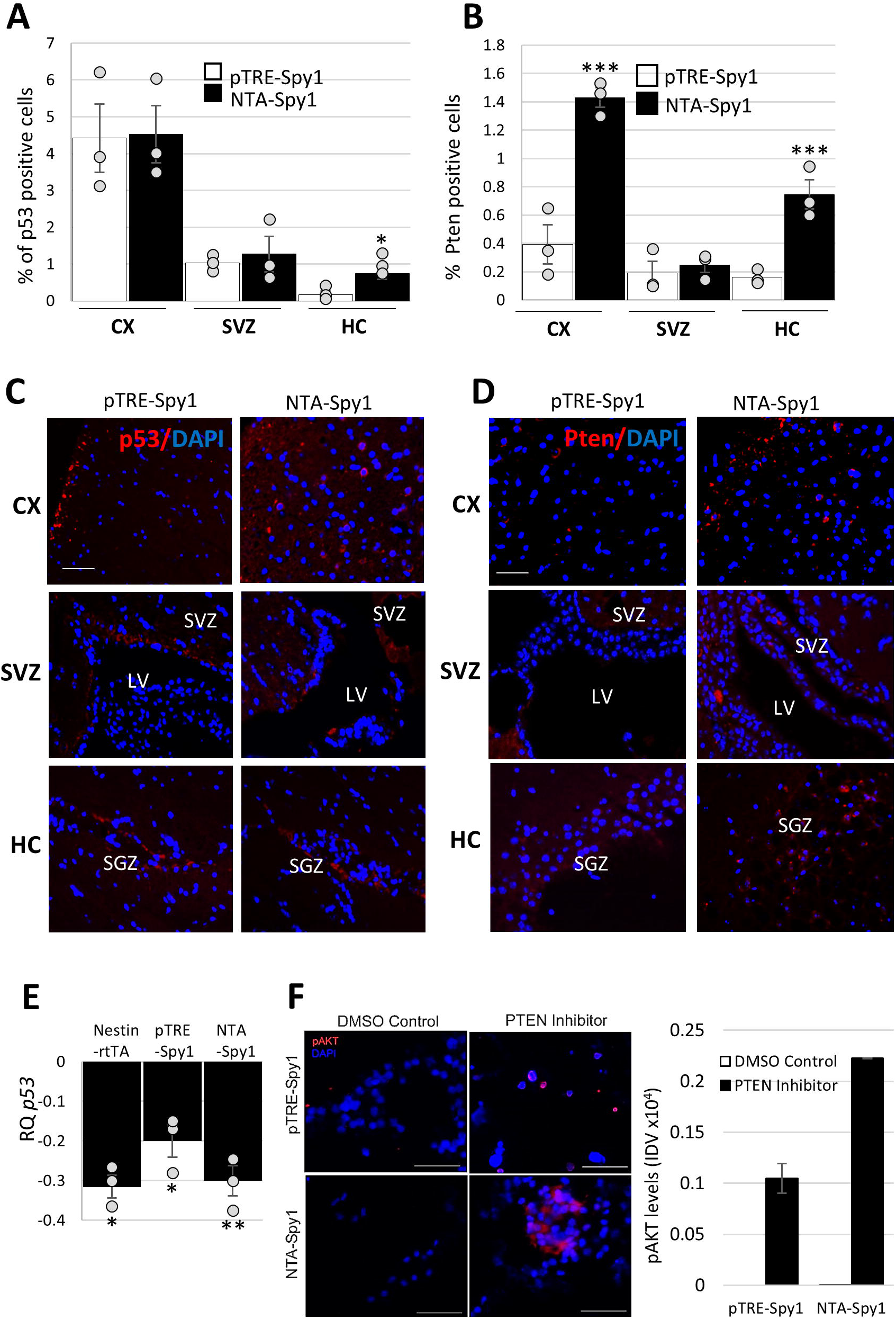

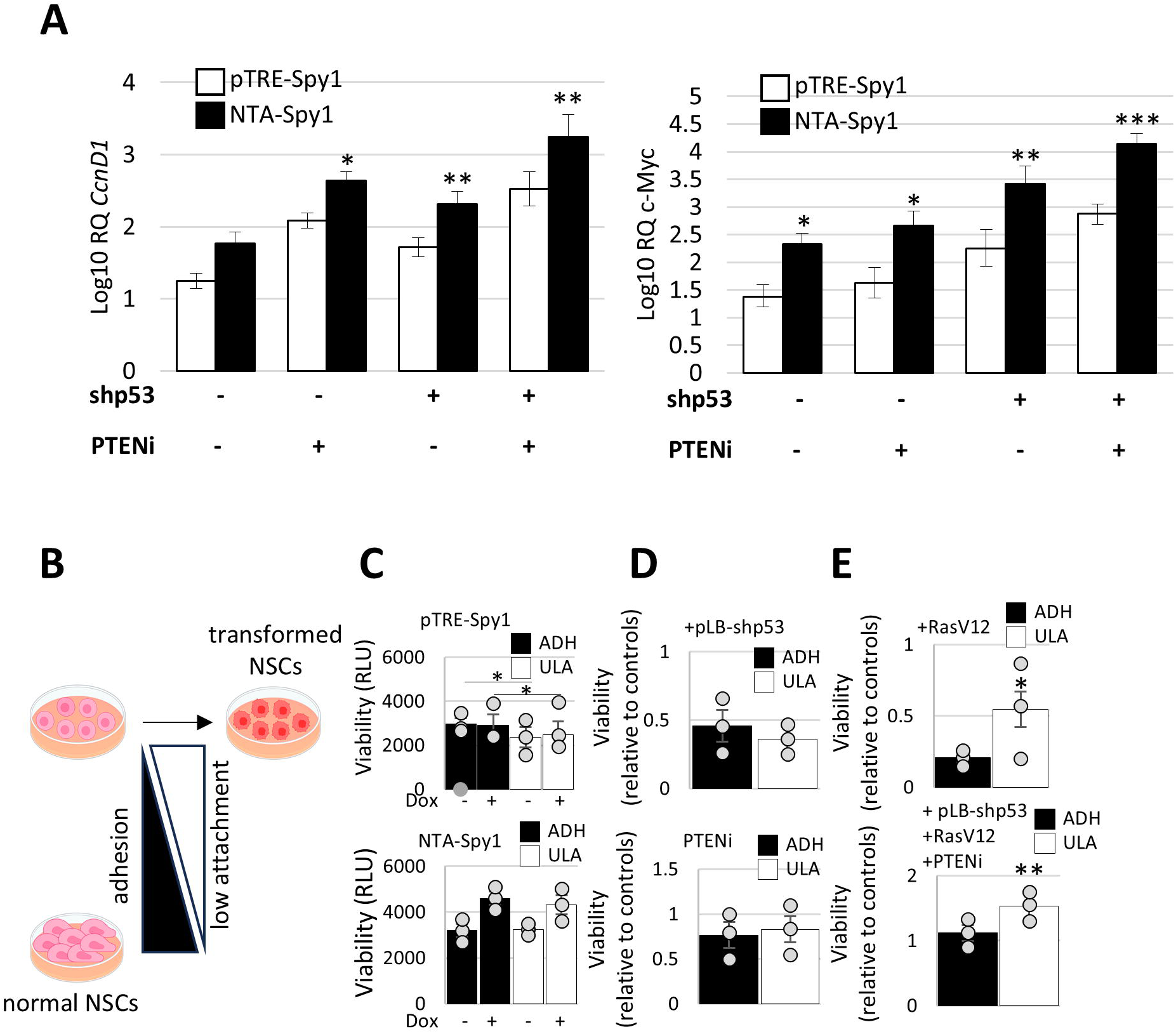

