## Supplemental information for "Atypical Cell Cycle Regulation over Neural Stem Cell Expansion"

**SUPPLEMENTAL MATERIAL**

**Supplemental Methods**

**Method Details**

**Generation of Spy1-pTRE Transgene**

Flag-Spy1 coding sequence was obtained from Flag-Spy1A-pLXSN ^55^. The pTRE-Tight Caspase 3 (p12):n2[TU#817] plasmid (Addgene 16084) was digested with EcoRI and Xbal to remove the Caspase 3. Flag-Spy1 coding sequence was then ligated into the pTRE-Tight backbone. Flag-Spy1-pTRE transgene was removed from the Spy1-pTRE-Tight vector by XhoI restriction digest.

**Generation of NTA-Spy1 Mice.**

The transgenic fragment was sent to London Regional Transgenic and Gene Targeting Facility to use for pronuclear injections into B6CBAF1/J hybrid embryos. PCR analysis was used to identify founders and maintain the colony. A 25μL PCR reaction was prepared comprising of 100-150ng of genomic DNA, 0.4μM forward primer, 0.4μM reverse primer (Supplemental Table 1), and 12.5μL of New England Biolabs Master Mix. The PCR cycling conditions used are as follows: 95°C for 2 minutes and 30 seconds; 95°C for 30 seconds, 60°C for 45 seconds and 72°C for 1 minute and 30 seconds (40 cycles); this was followed by a final extension at 72°C for 10 minutes.

Nestin-rtTA mice were purchased from the Jackson Laboratory [FVB/N-Tg (Nes-rtTA) 306Rsv/J] or from in house breeding at the University of Windsor. Genotyping was performed using 25μL PCR reactions containing 1 μL of genomic DNA, 0.4μM forward primer, 0.4μM reverse primer (Supplemental Table 1), and 12.5mL of New England Biolabs Master Mix. The PCR cycling conditions used are as follows: 94°C for 3 minutes; 94°C for 30 seconds, 57°C for 1 minute and 72°C for 1 minute (35 cycles); this was followed by a final extension at 72°C for 2 minutes. The Nestin-rtTA mice were crossed with pTRE-Spy1 mice to produce the Nestin-Spy1 (NTA) mice which were on a mixed genetic background (B6CBAF1/J FVB).

**Maintenance of NTA-Spy1 Mice.**

All animal experiments were conducted in accordance with Canadian Council on Animal care and the institutional guidelines for animal care and use. The protocol (AUPP #22-09) was approved by the University of Windsor Animal Care Committee. Animals were housed in a temperature-controlled facility maintained at 22 ± 2°C with a 12-hour light/dark cycle. Standard chow and water were provided ad libitum, and environmental enrichment was used to promote animal well-being. Mice were group-housed in ventilated cages with bedding material, with a maximum of three animals per cage, unless otherwise required by experimental conditions. Health checks were performed daily to monitor for any signs of distress or illness, and animals were acclimatized to their housing environment for at least one week prior to the initiation of experiments.

**Primary Neural Stem Cell Isolation**

Each mouse was humanely euthanized, and a midline incision was made in the skin along the length of the mouse. The skull was cut circumferentially with dissecting scissors and removed using forceps. Cranial nerves and blood vessels were cut, and forceps used to extract the brain, which was transferred to a 60 mm dish containing media with 3% penicillin/streptomycin (P/S) solution (Thermo Fisher Scientific, #15140122). The brain was cut in the coronal plane using a scalpel blade, and the area around the lateral ventricles isolated with visualization through a microscope. Isolated tissue was transferred to a sperate 60mm dish with 3% P/S solution (Thermo Fisher Scientific, #15140122), thoroughly minced with a scalpel blade and collected via centrifugation at 125g for 2 minutes. Supernatant was removed and 1 mL of Trypsin-EDTA (0.05%) (Thermo Fisher Scientific, #25300054) added. Suspension was incubated for 1 minute at 37°C before 1 mL of media containing 10% fetal bovine serum (FBS) (Thermo Fisher Scientific, #10437028) was added followed by incubation for 5 minutes at 37°C, subsequent washes with serum free media and collection followed by plating of the cells.

Subculture of primary cells

Cells were used for experiments within 7 days post-extraction by neurosphere collection via centrifuging at 500g for 5 minutes and dissociation of spheres using 1mL of Trypsin-EDTA (0.05%) (Thermo, #25300062) solution for 5 minutes at 37°C, followed by addition of 1mL of Neurobasal media containing 10% Fetal Bovine Serum (Thermo Fisher Scientific, #10437029) and manual dissociation.

**Neurosphere formation and clonal assays**

Neurosphere formation assays were conducted growth media in 6-well ULAPs (Corning™, #3471) at a density of 50,000 cells per well. Clonal assays were performed in 96-well ULAPs (Corning™, #07-201-680) in growth media. Cells were plated using serial dilution and wells containing 1-5 cells were included for experimental analysis.

**Differentiation assays**

For differentiation experiments, single cells were dissociated from primary neurospheres using PBS (Thermo Fisher Scientific, #10010023) and seeded in media supplemented with 2% FBS (Thermo Fisher Scientific, #10437029) as previously described^38^. In all lineage-specific differentiation protocols, single cells were seeded in poly-L-ornithine-coated (Sigma-Aldrich, #A-004-M) plates and supplemented with different growth mediums. For neuronal-specific differentiation, Neurobasal™ medium (Thermo Fisher Scientific, #21-103-049) supplemented with B-27™ supplement (Thermo Fisher Scientific, #17504044) and 2 mM GlutaMAX™ supplement (Thermo Fisher Scientific, #35050061) was added for 7 days. Dibutyryl cAMP (Sigma-Aldrich, #D0627) was added to the media with a final concentration of 0.5 mM and provided for 3 days. For astrocyte-specific differentiation, Dulbecco’s Modified Eagle Medium (DMEM) (Thermo Fisher Scientific, #11965092) supplemented with 1% N2 supplement (Thermo Fisher Scientific, #17502048), 2mM GlutaMAX™ supplement (Thermo Fisher Scientific, #35050061) and 1% FBS (Thermo Fisher Scientific, #10437029) was added for 7 days with a complete change of media every other day. For oligodendrocyte-specific differentiation, Neurobasal™ medium (Thermo Fisher Scientific, #21-103-049) supplemented with B-27 supplement (Thermo Fisher Scientific, #17504044), 2mM GlutaMAX™ supplement (Thermo Fisher Scientific, #35050061) and T3 (30 ng/mL; Sigma-Aldrich, #T6397) was added for 7 days with a complete change of media every other day.

**Matrigel Culture Assay**

Matrigel was supplemented with 20% growth media containing cells to obtain concentration of 1000 cells/20 μl of Matrigel. 20μl droplets of the mixture were pipetted onto the bottom of 6-well plate and incubated for 10 minutes at 37°C. 2mL of growth media were added to each well with the Matrigel droplets. Cell growth was monitored every 24 hours for 7 days and 3D images were taken via Z-stacking mode every 48 hours and analyzed. Cell colonies, equal to and/or larger than 50μm, were scored and cells with the neurite outgrowth double the length of cell body were counted as differentiated. Remaining cells which were not forming colonies or were not scored as differentiated were counted as non-differentiated cells. All cells were scored over the total number of cells seeded per Matrigel droplet.

**Quantitative Real Time-PCR (qRT-PCR) Analysis**

Cells used for RNA extraction were pelleted by centrifuging at 10,000 rpm. Total RNA was isolated using the RNeasy Plus Mini Kit (Qiagen, #74134) as per the manufacturer’s instructions. Subsequent cDNA synthesis was performed using QScript™ cDNA SuperMix (QuantaBio, #85048) as per the reagent’s accompanying manual. qRT-PCR reactions were then prepared using Fast SYBR™ Green Master Mix (Thermo Fisher Scientific, #4385612) as per the instructions in the associated manual and ran and analyzed using ViiA 7 Real-Time PCR System and associated software.

**Bromodeoxyuridine (BrdU) Proliferation Assay**

Cells were incubated in media containing BrdU (BD Biosciences, #550891) at a concentration of 10 μM overnight at 37°C and 5% CO_2_. Cells were subsequently fixed in 4% paraformaldehyde (PFA) solution in 1X phosphate-buffered saline (PBS) (Santa Cruz Biotechnology, #sc-281692) for 10 minutes at room temperature before incubation in 2 M hydrochloric acid (HCl) at 37°C for 20 minutes. Cells were then incubated with mouse anti-BrdU (BD Biosciences, #555627) in 0.1% Tween-20 (ACP Chemicals, #T-9780) in 1X PBS at room temperature for 45 minutes. Secondary antibody incubation was performed using 1:10 000 dilution of Goat Anti-Mouse IgG (H+L) Cross-Adsorbed Secondary Antibody Alexa Fluor™ 568 (Thermo Fisher Scientific, #A-11004) at room temperature for 1 hour. This was followed by Hoechst 33342 (Thermo Fisher Scientific, #H3570) counterstain at room temperature for 10 minutes. Imaging was performed using Leica CTR 6500 microscope.

**Immunocytochemistry**

Cells adhered to coverslips were washed with 1X PBS and fixed in 4% PFA in 1X PBS for 10 minutes at room temperature. Cells were subsequently permeabilized using 0.2% Triton X-100 (Biobasic, #TB0198) in 1X PBS at room temperature for 10 minutes before blocking with 10% normal goat serum (NGS) (Jackson ImmunoResearch, #005-000-121). Primary antibody incubation was performed using Anti-Phospho-Histone H2A.X (Ser 139) (Sigma-Aldrich, #05-636) at room temperature for 1 hour. Secondary antibody incubation was performed using 1:10000 dilution of Goat Anti-Mouse IgG (H+L) Cross-Adsorbed Secondary Antibody Alexa Fluor™ 568 (Thermo Fisher Scientific, #A-11004) at room temperature for 45 minutes before counterstain with Hoechst 33342 (Thermo Fisher Scientific, #H3570) to room temperature for 10 minutes. Coverslips were mounted on microscope slides and imaging was performed using Leica CTR 6500 microscope.

**Beta-III Tubulin Immunocytochemistry**

Cells adhered to coverslips were washed with 1X PBS and fixed in 4% PFA in 1X PBS for 10 minutes at room temperature. Cells were subsequently permeabilized using 0.2% Triton X-100 (Biobasic, #TB0198) in 1X PBS at room temperature for 10 minutes before blocking with 10% normal goat serum (NGS) (Jackson ImmunoResearch, #005-000-121). Primary antibody incubation was performed using anti-beta-III-Tubulin (MAB1195-SP) at 37°C for 2 hours at dilution 1:20 followed by washes with PBS, 3x 5 minutes and 1 hour incubation with Goat Anti-Mouse IgG (H+L) Secondary Antibody Alexa Fluor^TM^ 488 (Thermo Fisher Scientific #A-10670). Nuclei were counterstained with Hoechst. The fluorescent signal was scored across 3 fields of view as quantified as Integrated Density Values using ImageJ and divided by the total number of cells observed per field of view.

**Protein Isolation and Immunoblotting**

Cells were lysed with TNE buffer (50 mM Tris, 150 mM NaCl, 5 mM EDTA) for 20 minutes and supplemented with the following protease inhibitors: 2 μg/mL leupeptin (BioBasic, #LDJ691), 5 ug/mL aprotinin (Biobasic, #AD0153), 100 ug/mL PMSF (Biobasic, #PB0425). Cells were subsequently centrifuged at 4°C and 13 000 RPM for 15 minutes and the supernatant was collected. Protein concentration was measured using Bradford reagent (Sigma-Aldrich, #B6916) as per the instructions in the associated manual.

100 ug of protein per well was loaded on a 10% SDS (Biobasic, #SB0485) polyacrylamide (Biobasic, #A0007) gel and ran at 120 V for 3 hours. Gel was subsequently transferred to polyvinylidine fluoride (PVDF) membrane (Sigma-Aldrich, #IPVH00010) and ran at 30 V for 2 hours using a wet transfer method. Blocking of the membrane was performed at room temperature for 1 hour in 2% albumin fraction V (BSA) (Biobasic, #9048-46-8) or 2% milk (Thermo Fisher Scientific, #LP0033B), depending on the primary antibody used and the associated instructions. Primary antibody incubation was performed at 4 °C overnight. After primary antibody incubation, membrane was washed with Tris-Buffered-Saline-Tween 20 (ACP Chemicals, #T-9780) (TBST) 3 times for 8 minutes each time. Secondary antibody incubation was subsequently performed at room temperature for 1 hour, followed by 3 washes of 5 minutes each with TBST. Chemiluminescent peroxidase substrate (Thermo Fisher Scientific, #32106) was used for imaging of the membrane as per the instructions in the associated manual. Imaging was performed using FluorChem® HD2 Gel Imaging System and densitometry analysis was performed with AlphaEaseFC software.

Primary antibodies used and their respective dilutions were as follows: Anti-Actin, 1:1 000 (Sigma-Aldrich, #MAB1501); Anti-Spy1, 1:700 (Abcam, #ab220000); Anti-Sox2, 1:1 000 (Cell Signalling Technology, #2748); Mouse Anti-Klf4, 1:800 (R&D Systems, #AF3158), Anti-p27 KIP 1 (Abcam, #ab137736); Anti-Musashi 1 (Msi1), 1:1 000 (Abcam, #ab21628); Anti-NUMB, 1:800 (Abcam, #ab4147).

Secondary antibodies used and their respective dilutions are as follows: Normal Mouse IgG, 1:10 000 (Sigma-Aldrich, #12-371); Normal Rabbit IgG, 1:10 000 (Sigma-Aldrich, #12-370).

**Caspase 3/7 Cell Death Assay**

The Caspase-Glo**®** 3/7 Assay System (Promega Corporation, #G8091) was used in accordance with the instructions outlined in the associated manual. Luminescence was measured using a SpectraMax (Molecular Devices).

**Flow cytometry and Fluorescence-Activated Cell Sorting (FACS) of Nestin+ cells**

NSCs were isolated and cultured as neurospheres were dissociated using PBS (Thermo Fisher Scientific, #10010023) and the suspension was washed through a 70 µm strainer (Grenier, #542070). Cells were then counted using TC-10 automated cell counter (Bio-Rad, #145-0001) and resuspended in ice-cold PBS (Thermo Fisher Scientific, #10010023) at a maximal concentration of 1 x 10^6^ cells. Cells were fixed using fixation buffer (BD Biosciences, #554655), permeabilized with permeabilization/wash buffer (BD Biosciences, #557885) and stained using stain buffer supplemented with FBS (BD Biosciences, #554656). Cells were resuspended at a maximal concentration of 1 x 10^6^ cells/mL of stain buffer and stained using mouse/rat Nestin PE-conjugated antibody (R&D Systems, #IC2736P, 10 μL) or CD24-APC conjugated antibody (BD Biosciences, #562349, 1.7 μL/50 μL test) for direct staining. Cells were stained in the dark on ice for 1 hour, and vortexed every 10 minutes. After 1 hour, cells were spun down at 500g for 5 minutes and staining solution was removed. Anti-EGFR primary antibody (Cell Signaling, #4267T, 0.25μL/50 μL) and anti-Sox2 antibody (Abcam, #ab97959, at 5μg/mL) were incubated with cells at concentration 1 x 10^6^ cells/mL in the stain buffer for 2 hours at room temperature, washed with 1XPBS and incubated with secondary, fluorophore conjugated antibody (BD Bioscienecs, 1:1000). Cells were washed twice in wash buffer (BD Biosciences, #557885) and resuspended in ice-cold PBS (Thermo Fisher Scientific, #10010023) for FACS. All flow cytometry experiments were performed using BD LSRFortessa X-20 with BD High Throughput Sampler (HTS), FACSAria with FACSDiva software. Cell debris was excluded from the final analysis. Sorted cells were pelleted via centrifuging at 10,000 rpm and subjected to cell lysis and RNA extraction using RNeasy Plus Mini Kit (Qiagen, #74134) followed by cDNA synthesis and qRT-PCR.

**Transfection of Primary NSCSs**

pTRE-Spy1 and NTA-Spy1 neurospheres were dissociated into single cell suspensions using PBS (Thermo Fisher Scientific, #10010023) and plated on 0.1% gelatin-coated 6 cm plates and allowed to adhere overnight. The Lipofectamine 2000 (Thermo Fisher Scientific, #11668019)/mouse *Msi*1 siRNA (Thermo Fisher Scientific, #AM16708) complex was prepared in 400 μL of media and allowed to incubate at room temperature for 15 minutes. The final volume of Lipofectamine 2000 added per plate was 12 μL and the final mass of siRNA added to each plate was 1.5 μg. The transfection media was added to the plates in a dropwise manner. Cells were incubated in transfection media for 5 hours after which the transfection media was removed, and normal growth media added. Cells were used for experiments for up to 48 hours post-transfection.

**Growth in Low Attachment Conditions (GILA) Assay**

GILA assay was conducted as described before^32^. 1000 cells/well were seeded into 96-well plates with high attachment surface (Sarstedt, #83.3924) or ultra-low attachment (Corning™, #07-201-680) in 100 μL of growth media as outlined above and cultured for 5 days. Cell viability was then assayed using CellTiter-Glo® 3D Cell Viability Assay (Promega Corporation, #G9681) as per the instructions in the associated manual and a plate reader. Luminescence was quantified as Relative Luminescence Units (RLUs) and values of the treatment cohort were corrected by values of the respective controls. Viability was compared between high attachment and ultra-low attachment conditions and graphed as a ratio.

**Immunostaining Brain Tissues**

Intact brain tissue was removed from mice as described above and placed in 10% neutral buffered formalin. Intact brain tissue was dehydrated through a series of ethanol solutions of increasing concentration and cleared in xylene. The tissue was then embedded in paraffin wax (*) and sectioned into 5 μm sections using Leica RM2125RT microtome. Sections were rehydrated through a series of ethanol solutions of decreasing concentration and heat-mediated antigen retrieval was performed using a 10 mM sodium citrate buffer or EDTA buffer depending on the antibody manufacturer’s recommendations. Sections were subsequently blocked at room temperature for 1 hour using 3% BSA (Biobasic, #9048-46-8)/0.1% Tween 20 (ACP Chemicals, #T-9780) in 1X PBS before incubation with anti-Nestin (Biotechne, #NB100-1604), anti-EAAT1 primary antibody (Cell Signaling, #5684T), anti-EGFR primary antibody (Cell Signaling, #4267T), anti-PCNA primary antibody PC10 (Santa Cruz, #sc56) in blocking solution (1:200 dilution), or Sox2 antibody (Abcam #97959; at 1μg/mL) or Spy1 antibody (Abcam # ab153965, at 1μg/mL), Anti-NeuN antibody[EPR12763] (Abcam#177487; at 1:1000), Brn2/POU3F2 (D2C1L) antibody (Cell Signaling #12137; at 1:500) and anti-Neurofilament heavy polypeptide antibody [EPR20020] (Abcam #207176; at 1:1000) at 4°C overnight. Sections were then incubated with [goat anti-Rabbit or anti-Mouse IgG (H+L) Cross-Adsorbed Secondary Antibody, Alexa Fluor™ 488](https://www.thermofisher.com/antibody/product/Goat-anti-Rabbit-IgG-H-L-Cross-Adsorbed-Secondary-Antibody-Polyclonal/A-11008) (Invitrogen #A-11008 or #A-11001, respectively) and/or Alexa Fluor™ 568 (Invitrogen #A-11011-rabbit or #A-11004- mouse) at 1:1000 dilution at room temperature for 1 hour in a humidified chamber. Sections were mounted using Permount™ mounting medium (Fisher Chemical™, #SP15-100) and imaged with Leica CTR 6500 microscope. Regions of the coronal sections targeted for the analysis of the fluorescent signal were identified per Allen Mouse Brain Atlas (atlas.brain-map.org, Allen Institute). For the analysis of the cortical layers, the same slice position was assured across all samples and controls analyzed. ImageJ software was used to score the fluorescent signal.

**Orthotopic Cell Transplantation into Mice**

pLB-shSpy1- transduced GL261 cells along with the control were selected based on EGFP expression of the pLB vector via FACS (FACSAria) prior to transplantation and 100% transduced cells were used. On postnatal day 1, C57BL/6 mice were cryoanaesthetized and 25 000 GL261 cells were orthotopically transplanted into the brain using stereotactic injector (Stoelting Co.). The orthotopic injection was guided using Neurostar software (*) with the bregma as the zero point and coordinating the injection site to X, Y, Z = (0.8, 2.0, -1.5 mm). The cells were injected in a total volume of 2 μL at a rate of 30 nL/min over 5 min and the needle was left in place for an additional 2 minutes to prevent reflux. The mice were monitored during recovery. Tumor growth and progression were assessed using SII Ami-HT *in vivo* imager and fluorescence intensity was measured using Aura software and quantified as Average Radiant Efficiency over 8 mice per treatment.

**Beta-Galactosidase (β-Gal) Stain**

Cells were fixed in 4% PFA (Santa Cruz Biotechnology, #sc-281692) at room temperature for 10 minutes. Staining solution (5 mM K_4_[Fe(CN)_6_]·3H_2_O, 5 mM K_3_[Fe(CN)_6_] and 2 mM MgCl_2_) was prepared in 1X PBS. 25 uL of 40 mg/mL X-Gal (Biobasic, #BB0083) was added to every mL of staining solution to generate complete staining solution, and this was performed directly before staining. Cells were stained at 37°C overnight and subsequently imaged using the Leica CTR 6500 microscope. To quantify SA-β-Gal activity, ImageJ software was used to determine the percentage area of positive staining. Colour deconvolution was performed using the Alcian Blue & H vector to separate the colour components of the staining within the image. A threshold was adjusted to isolate the positive area of interest, and the corresponding percentage area (% Area) was measured.

**Immunofluorescent Cell Pair Assay**

Single cells were isolated using 0.05% Trypsin-EDTA (Thermo Fisher Scientific, #25300054) and plated onto MaxGel™ ECM (Sigma-Aldrich, #E0282)-coated coverslips. Cell division was monitored over 20 hours and cells were then fixed using 4% PFA (Santa Cruz Biotechnology, #sc-281692) at room temperature for 15 minutes. Cells were then washed with 1X PBS, permeabilized with 0.2% Triton X-100 (Biobasic, #TB0198) in 1X PBS and incubated with anti-NUMB primary antibody (Cell Signaling #C29G11; 1:400 dilution) or anti-NICD antibody (Cell Signaling #4147T; 1:200 dilution) or CD133 antibody (Thermo Fisher Scientific # PA538014) for 2 hours at 37°C. Cells were then incubated with Goat Anti-Rabbit IgG (H+L) Cross-Adsorbed Secondary Antibody Alexa Fluor^TM^ 488 (Thermo Fisher Scientific, #A-11008) for NUMB and Goat Anti-Rabbit IgG (H+L) Cross-Adsorbed Secondary Antibody Alexa Fluor^TM^ 568 (Thermo Fisher Scientific, #A-11011) for NICD and CD133, for 1 hour at room temperature. TOTO-3 (Thermo Fisher Scientific, #32106) for NUMB and Hoechst (for NUCD and CD133) were used as a nuclear counterstain. Five random fields of view were scored for the tested protein distribution in mitotic cell pairs/treatment/replicate. Imaging and analysis were performed using Leica CTR 6500 microscope and Leica LAS X software, respectively.

**Novel Object Recognition Test**

Novel object recognition test was performed as described^71^. Plexiglass boxes with dimensions of 40 cm x 40 cm x 40 cm were built and covered with frosted white paper to impair outside views from inside the box. Plastic cubes that were either blue or yellow with dimensions of 4 cm x 4 cm x 4 cm were used as objects. Computer cameras were used to record videos. On days 1 and 2, mice were moved to an area near the boxes for 10 minutes. To acclimate the mice to their environment and decrease stress, the mice were then also individually placed directly in the boxes for 10 minutes on both days. Plexiglass boxes and the objects were cleaned with 70% ethanol after every mouse and dried for 10 minutes before the next mouse was placed inside. On day 3, mice were transported to an area near the boxes and allowed to sit in cages for 10 minutes. As before, mice were then also placed in boxes to explore for 10 minutes and were recorded. On day 4 testing was performed. One of the old objects was replaced with a novel object. Similarly, mice were taken to an area near the boxes, allowed to sit for 10 minutes and then placed inside the boxes and recorded for 10 minutes. The exploration time was counted by the period that the mice stayed facing the object within a 5 cm radius of the object. Exploration times of under 20 seconds or over 5 minutes at an object were excluded. Preference rate was measured by dividing the total time spent exploring both objects by the time spent exploring one object.

**Lentiviral Constructs, Production and Transduction**

Lentiviral vectors pEIZ-flag-SPY1, pLB-shSpy, pLB-shp53, pLENTI-CMV-RasV12, along with control vectors were used to generate lentiviral particles by co-transfecting HEK293T cells with the expression vector, along with the packaging plasmids (pMD2.G, pRSV-Rev, pMDLg/pRRE). Viral supernatants were collected 48 hours and 72 hours post-transfection, filtered, and concentrated by ultracentrifugation for 4 hours at 24,000 rpm. Primary cultures collected from SVZ of P2 mice were transduced with lentiviral particles in the presence of 8 µg/mL polybrene at MOI=5. Cells expressing ZsGreen (pEIZ) or EGFP (pLB) post infection were selected/sorted using FACS (FACSAria) to generate lines which 100% of population is modified for the gene or shRNA expression. Successful transduction was validated by qRT-PCR assessment of gene expression levels vs. control.

**Zebrafish Xenografts**

Zebrafish handling and xenograft experiments were performed following the guidelines approved by University of Windsor Institutional Animal Care and Use Committee (AUPP#22-01). Zebrafish were housed at 28^o^C; 50 fish per tank. Cells were labelled with fluorescent DiO (green) or DiI (red) dye to track the cells post injection. 72 hours post fertilization (hpf) 9nL of cell suspension of 1*10^6^ cells/mL were injected into the zebrafish yolk sac using Nanoject III Injector (Drummond Scientific Company; CA#3-000-207) with a stereo microscope. After injection, the embryos were transferred into 33^o^C. Xenograft growth and dissemination were monitored by fluorescence microscopy and images were taken for analysis using ImageJ.

**Transfection of Primary NSCs**

pTRE-Spy1 and NTA-Spy1 neurospheres were single cell dissociated and plated on 0.1% gelatin coated 6cm plates and allowed to adhere overnight. The Lipofectamine 2000 (Thermo)/siRNA (mouse si Msi1; Thermo) complex was prepared in 400µL media and allowed to incubate at room temperature for 15 minutes. Final Lipofectamine 2000 reagent per plate used was 12µL and the final amount of siRNA added to each plate was 1.5µg. The Lipofectamine/siRNA transfection media was added dropwise to plates. Cells were incubated in transfection media for 5 hours. This was followed by a media change and cell were used for experiments 48 hours post transfection.

**Caspase 3/7 Cell Death Assay**

The Caspase-Glo 3/7 Assay (Promega) was done as per manufacturer’s instructions. Luminescence was measured through a luminometer.

**Nestin Fluorescence Associate Cell Sort (FACS)**

NSCs were harvested, and neurospheres dissociated. Cells were washed through a 70µm cell strainer and collected at a concentration of 1 x 10^6^ cells/mL of ice-cold PBS. To aid in intracellular staining for Nestin, cells were fixed with fixation buffer (BD Pharmingen) and permeabilized with permeabilization/wash buffer. Cells were stained using BD Pharmingen Stain Buffer containing FBS. 1 x 10^6^ cells/mL of buffer. Anti-mouse/rat Nestin PE conjugated antibody (R&D Systems) was used at a concentration of 10µL/mL of stain buffer. Cells were stained in the dark on ice for 1 hour, with vortexing every 10 minutes. Cells were spun down, and staining solution was removed. Cells were washed twice in wash buffer and resuspended in ice cold PBS for FACS. All flow cytometry experiments were performed using BD LSRFortessa X-20 with FACSDiva software. Cell debris was excluded from analysis.

**Growth In Low Attachment Conditions (GILA) Assay**

GILA assay was conducted as described before ^32^. 1000 cells/ well were seeded into 96 well plates of high attachment surface (Sarstedt) and/or Ultra Low Attachment 96 well plates (Greiner) in 100ul of Neurobasal growth media and cultured for 5 days. Cell viability was then assayed using a plate reader and 3D Cell Titer Glo (Promega). Luminescence was quantified as Relative Luminescence Units (RLU) and the values in the treatment cohort were corrected by values of respective controls. Viability was compared between high attachment and Ultra Low Attachment conditions and graphed as their ratio, respectively.

**Immunostaining of brain tissues**

Brain tissue was collected and fixed in 10% neutral buffered formalin. Brain tissue was dehydrated through a series of increasing concentrations of ethanol and cleared in xylene. Tissues were embedded in paraffin wax and sectioned into 5µM sections using Leica RM2125RT. Sections were rehydrated through a series of decreasing ethanol concentrations and heat mediated antigen retrieval was performed with 10mM sodium citrate buffer. Sections were blocked at room temperature for 1 hour with 3% BSA- 0.1% Tween-20 in 1X PBS for rabbit secondary antibodies. Sections were incubated with primary PCNA (1:200) antibody in blocking solution overnight at 4°C. Sections were incubated in secondary antibody (1:750) for 1 hour at room temperature in a humidified chamber. Sections were mounted using Permount toluene solution (Thermo) and imaged with Leica CTR 6500 microscope (Leica Microsystems, Heidelberg, Germany).

**Statistical Analyses**

All results are presented as mean values ±SEM.

Statistical analyses were performed using GraphPad Prism 10.4.1 software using Student’s *t*-test, one-way and two-way ANOVA.

Student’s *t*-test was used and a p-value of less than 0.05 was considered significant. All data are reported as mean ± standard error of the mean (SEM).

For qRT-PCR analysis, cycle threshold (Ct) values of the internal control gene (*GAPDH*) were subtracted from the corresponding Ct value of a target gene which generated a ΔCt value which was then subjected to the Student’s *t*-test analysis.

For comparative analysis between (WT), Nestin-rtTA, pTRE-Spy1 and NTA-Spy1 mice and primary cultures, littermate mice were used.

For neurosphere size analysis one-way ANOVA was used with Post Hoc Tukey HSD for pairwise comparisons between NTA-Spy1 and controls; ***p < 0.0001; ****p < 0.00001. Shown on the graphs is comparison between NTA-Spy1 and pTRE-Spy1 derived spheres.

**Supplementary tables**

| **Gene** | **Forward sequence (5’-3’)** | **Reverse sequence (3’-5’)** |
| --- | --- | --- |
| *Gapdh* | GATGCCCCCATGTTTGTGAT | GTGGTCATGAGCCCTTCCA |
| *Spy1* | TCATACAGCGCCAGGAAATG | AAGGTCATAGCCAAAAGATACTTGTCT |
| *Flag* | TGACAAGAGGCACAATCAGATGT | CAAATAGGACGCTTCAGAGTAATGG |
| *CD133* | GAAAAGTTGCTCTGCGAACC | CTCGACCTCTTTTGCAATCC |
| *Sox2* | GGGAAATGGGAGGGGTGCAAAAGAGG | TTGCGTGAGTGTGGATGGGATTGGTG |
| *Nestin* | ACCTATGTCTGAGGCTCCCTATCCTA | ACCTATGTCTGAGGCTCCCTATCCTA |
| *Oct4* | ATGGCATACTGTGGACCTCA | AGCAGCTTGGCAAACTGTTC |
| *Klf4* | ACGATCGTGGCCCCGGAAAAGGACC | TGATTGTAGTGCTTTCTGGCTGGGCTCC |
| *c-Myc* | GCGTCCTGGGAAGGGAGATCCGGAGC | TTGAGGGGCATCGTCGCGGGAGGCTG |
| *Nanog* | CTCATCAATGCCTGCAGTTTTTCA | CTCCTCAGGGCCCTTGTCAGC |
| *S100 Beta* | TGGCCACCAGTAACATGCAA | CAGTTGGCGGCGATAGTCAT |
| *O4* | CTTCCTCGACCAGACCTCG | ACAGGATCGGTTCGGAGTGT |
| *Mapt* | CGCTGGGCATGTGACTCAA | TTTCTTCTCGTCATTTCCTGTCC |
| *Beta III Tubulin* | TAGACCCCAGCGGCAACTAT | GTTCCAGGTTCCAAGTCCACC |
| *Msi1* | TAGTTCGAGGGACAGGCTCT | GTTGAGGGACAGGCAGTAGC |
| *Hes1* | TGAAGGATTCCAAAAATAAAATTCTCTGGG | CGCCTCTTCTCCATGATAGGCTTTGATGAC |
| *Hes5* | TGCAGGAGGCGGTACAGTTC | GCTGGAAGTGGTAAAGCAGCTT |
| *Vimentin* | CAGCAGTATGAAAGCGTGG | GGAAGAAAAGGTTGGCAGAG |
| *Bmi1* | TATAACTGATGATGAGATAATAAGC | CTGGAAAGTATTGGGTATGTC |
| *Ki67* | TGCAGAGAATGTCGGGATAAAG | AGGAGATGGAGAAGTGAAGAGG |
| *Cdkn1a (p21)* | CCTGGTGATGTCCGACCTG | CCATGAGCGCATCGCAATC |
| *Ink4a (p16)* | ATGGAGCCGGCGGCGGGGAGCAGCATGGAGCCT | TCAATCGGGGATGTCTGAGGGACCTTCCGC |
| *p53* | CACAGCGTGGTGGTACCTTATG | GGTTCCCACTGGAGTCTTCCA |
| *pTRE-Spy1 (Set 1)* | GTGTACGGTGGGAGGCCTATATAA | GTCATAGCCAAAAGATACTTGTCTGC |
| *pTRE-Spy1 (Set 2)* | GTATGTCGAGGTAGGCGTGT | CTATCCCAGAGCTGGTCCCT |
| *Nestin-rtTA Genotype* | CGCTGTGGGGCATTTTACTTTAG | CATGTCCAGATCGAAATCGTC |

**Supplemental Table 1. Primers used for both RT-PCR and qRT-PCR analysis.** The primers are listed with their name, their purpose (RT or qRT), the sequence of forward (5’🡪3’) primer and the sequence of reverse (5’🡪3’) primer.

**Supplementary figure legends.**

**Figure S1. Generation of pTRE-Spy1 transgenic mouse model.**

(A) Complete plasmid map illustrating pTRE-Spy1 vector.

(B) Agarose gel image depicting RT-PCR analysis of 536bp portion cut from vector in (A) using XhoI and AlwNI.

(C) Agarose gel image depicting RT-PCR of the 536bp band of founder pTRE-Spy1 mice. Yellow boxes indicate positive founders.

(D) Agarose gel image depicting RT-PCR corresponding to successful germline transmission of the pTRE-Spy1 transgene (red arrow). Numbers in (C) and (D) are indicative of the mouse tag number belonging to each tail sample that was screened. Vectors represents the pTRE-Spy1 plasmid used as a positive control.

(E) Representative gating of FACS assay of Nestin+ cells used in sorting of pTRE-Spy1 and NTA-Spy1 SVZ derived cells analyzed in Figure 1D.

(F) Flow cytometry analysis of Nestin+ cells (left) derived from a SVZ and propagated as neurospheres. Cells quantified as percentage of the population tested (right). Three separate cultures from an uninduced NTA-Spy1 mouse at PN2 analyzed. N=3; results presented as the mean ±s.d.; ***p < 0.001; Student’s *t*-test.

(G) Flow cytometry analysis of Sox2+ cells as a percentage of Nestin+ populations in (G).

(H) Percentage of GLAST + and – cells in CD24+/- populations derived in experiment shown in Figure 1E. N=3, results presented as the mean ±s.d.; ***p < 0.001, Student’s *t*-test.

(I) Analysis of the percentage of cells positive for GLAST (left) and EGFR (right) in NTA-Spy1 neurosphere cultures treated with 1μg/mL of doxycycline. SVZ from one individual PN2 NTA-Spy1 mouse was dissected and dissociated into three separate cultures. N=3, results presented as the mean ±s.d.; ***p < 0.001; Student’s *t*-test.

**Figure S2. Lentiviral overexpression of Spy1 increases stemness markers.**

qRT-PCR analysis of gene expression levels of Msi1, Vimentin, Bmi1, Nestin and CD133. Transient overexpression of Spy1 is achieved via lentivirus infection of either SPY1 (pEIZ-SPY) or an empty control vector (pEIZ-Cnt). Gene expression levels are presented as Relative Quantification (RQ) in mouse primary neurosphere cultures for pEIZ-SPY or pEIZ-Cnt**.** Data analyzed over three separate infections (n=3) and shown as mean ± s.d., *p < 0.05, **p < 0.01: Student’s *t*-test.

**Figure S3. NTA-Spy1 cells maintain self-renewal ability of differentiating NSCs.**

Primary cell cultures were extracted from SVZ of mice between PN day 2 and 4 of age. Cells were maintained and assayed *in vitro* in the presence of 1μg/ml of doxycycline.

(A) Western blot of Sox2 protein levels (top) in pTRE-Spy1 and NTA-Spy1 cells. Actin used as a loading control. Densitometry analysis (bottom) of Sox2/Actin IDV in pTRE-Spy1 and NTA-Spy1 cells. Cultures from three individual mice per genotype were used to prepare cell lysates. N=3, results presented as the mean ±s.d.; *p < 0.05; **p < 0.01. Student’s *t*-test.

(B) QRT-PCR analysis of Olig2, Sox2, Pouf3 and, Sall2 gene expression levels scored as Relative Quantification (RQ) in NTA-Spy1 cells corrected by pTRE-Spy1 control. RNA was extracted from three mice per genotype. N=3, results presented as the mean ±s.d.; *p < 0.05; Student’s *t*-test.

(C) Schematic showing experiment set-up for the re-sphering assay.

(D) Percentage of cells with the capacity to re-form a sphere quantified as spheres formed over the total number of pTRE-Spy1 and NTA-Spy1 cells seeded (150000 over 4 individual experiments/mice per genotype) (left). Representative light microscope images of pTRE-Spy1 and NTA-Spy1 cells depicting the “re-sphere” in the NTA-Spy1 culture (right). Scale bar = 50 µm. N=4, results presented as the mean ±s.d.; **p < 0.01; Student’s *t*-test.

(E) QRT-PCR analysis of differentiation markers S100β, O4, and Mapt in primary neurospheres (NS), differentiated cells and “re-sphering cells” of pTRE-Spy1 and NTA-Spy1- derived cultures from three mice per genotype. N=3, results presented as the mean ±s.d.; *p < 0.05, ***p < 0.001; Student’s *t*-test.

(F) Immunofluorescent staining of beta-III Tubulin in the absence and presence of doxycycline in in three separate cultures derived from an NTA-Spy1 mouse. DAPI used as a nuclear counterstain. Scale bar = 10 µm. N=3, results presented as the mean ±s.d.; ***p < 0.001; Student’s *t*-test.

**Figure S4. Distribution of cell fate- determinants depends on the expression of Spy1 and is regulated via Msi1 pathway.**

(A) Right: pTRE-Spy1 (top) and NTA-Spy1 (bottom) cells of three individual mice, obtained from SVZ regions of the brain at PN2, cultured in Matrigel^TM^ in the presence of 1μg/ml of doxycycline. Arrows point at morphology of colonies formed (red), differentiated cells (green) and non-differentiated cells (blue); scale bar = 75 µm. Right: Colonies formed, differentiated cells and non-differentiated cells quantified as percentage of the number of cells seeded. N=3, results presented as the mean ±s.d.; **p < 0.01; Student’s *t*-test.

(B) Cell pair assay in MaxGel ECM in the presence of 1μg/ml of doxycycline. Representative images of Numb protein distribution in mitotic pairs (left) of pTRE-Spy1 (top) and NTA-Spy1 (bottom). Mitotic cell pairs were scored (right) as asymmetric or symmetric and graphed as a percent of total mitotic pairs counted and averaged over three pTRE-Spy1 and three NTA-Spy1individual cultures. Scale bar = 5 µm. N=3, results presented as the mean ± s.d.; **p < 0.01; Student’s *t*-test.

(C) CD133 protein distribution in mitotic pairs of cell pair assay using NTA-Spy1 cells derived at PN2, in presence an absence of doxycycline.

(D) Schematic of Msi1 signaling.

(E) Western blot for Msi1, Numb (left). Actin used as a loading control. Densitometry analysis (right) of Musashi 1/Actin Integrated Density Values (IDV) and Numb/Actin IDV in three individual pTRE-Spy1 and three NTA-Spy1mouse- derived cells. N=3, results presented as the mean ± s.d.; **p < 0.01; Student’s *t*-test.

(F) QRT-PCR analysis of gene expression levels of Msi1 (left) and Hes1 and Hes5 (right) in cells from pTRE-Spy1 and NTA-Spy1mice and treated with siMsi1 and 1μg/ml of doxycycline. Expression quantified as LogRQ values relative to siControl. Three cultures from one mouse/genotype tested. N=3, results presented as the mean ± s.d.; *p < 0.05, ***p < 0.001; Student’s *t*-test.

(G) Cell pair assay in pTRE-Spy1 and NTA-Spy1 siMsi1 cells vs. control (siControl) graphed as percentage of cell pairs presenting symmetric distribution of Numb over total number of cell pairs scored. Four individual cultures per genotype tested. N=4, results presented as the mean ± s.d.; *p < 0.05, **p < 0.01; Student’s *t*-test.

(H) NICD distribution in mitotic pairs of the cell pair assay in NTA-Spy1 cells without (-doxycycline) or with doxycycline (+doxycycline) treatment. Representative images (left). Scale bar = 20 μm. Data graphed as percentage of pairs presenting symmetric vs. asymmetric distribution of NICD signal over total number of mitotic pairs in the assay. Three individual NTA-Spy1 cultures tested. N=3, results presented as the mean ±s.d.; **p < 0.01, ***p < 0.001; Student’s *t*-test.

**Figure S5. NTA-Spy1 mice demonstrate shift in distribution and density of cortical layers.**

Coronal sections obtained from matched regions of the brain of NTA-Spy1 and pTRE-Spy1 mice at 20 months of age, subjected to immunostaining with selected antibodies. Hoechst stain used to mark nuclei.

(A&B) Neurons positive for NeuN and Pou3f2 expression, respectively, scored and quantified over total number of nuclei per field of view in each cortical layer.

(C) Intensity of fluorescent immunostain against NFH quantified over nuclear signal per filed of view using ImageJ.

Fields of view in two separate cortical regions per section were analyzed in three individual animals per genotype. N=6, results presented as the mean ±s.d.; *p < 0.05, ***p < 0.001; Student’s *t*-test.

**Figure S6.** (A)&(B) Immunohistochemistry- based analysis of p53 (A) and (Pten) protein expression levels in cortex (CX), subventricular zone (SVZ) and, hippocampus (HC) of pTRE-Spy1 and NTA-Spy1 mice at 4 months of age. Number of positive cells scored and quantified over total cell number in a filed of view, over five fields of view in three separate mice, using ImageJ. DAPI used as a nuclear counterstain. N=3, results presented as the mean ±s.d.; *p < 0.05, ***p < 0.001; Student’s *t*-test.

(C)&(D) representative images of the CX (cortex), SVZ (subventricular zone), and HC (hippocampus) immunostained to assess levels of p53+ and Pten+ in Figure S4A&B. Neurogenic regions marked in the image; LV= Lateral Ventricles, SGZ= Subgranular zone.

(E) p53 knockdown confirmed using qRT-PCR in Nestin-rtTA, pTRE-Spy1 and NTA-Spy1 cells infected with pLB-scrambled control vector or pLB-shp53 in cultures derived from three mice per genotype. N=3, results presented as the mean ±s.d.; *p < 0.05, **p < 0.01; Student’s *t*-test. (F) Immunocytochemistry confirming the inhibition of PTEN activity by phosphorylated AKT (pAKT) levels in pTRE-Spy1 and NTA-Spy1 cells treated with PTEN inhibitor or control. pAKT fluorescent signal quantified using ImageJ and graphed as Integrated Density Values relative to nuclear signal.

**Figure S7.** **Spy1 overexpression induces aberrant changes in characteristics of normal NSCs**

(A) qRT-PCR analysis of mRNA expression levels of *CcnD1* and *c-Myc* in neurosphere cultures derived from littermate pTRE-Spy1 and NTA-Spy1 mice at 4 months of age and transduced with shp53 and/or treated with PTEN inhibitor (PTENi). Results presented as the mean ±s.d.; *p < 0.05, **p < 0.01, ***p < 0.001; Student’s *t*-test.

(B) Schema demonstrating growth preference in normal and transformed cells. (C-E) GILA (Growth In Low Attachment) assay in pTRE-Spy1 and NTA-Spy1 cells derived from three individual mice/genotype at 4 months of age cultured in ultra low attachment (ULA) conditions in comparison to high attachment (ADH) in the presence/absence of doxycycline. (C) Viability of pTRE-Spy1 (top) and NTA-Spy1 (bottom) cells. (D) NTA-Spy1 cells transduced with shp53 (pLB-shp53), top, or treated with PTENi (bottom) and induced with doxycycline, (E) NTA-Spy1 cells transduced with RasV12 (pLENTI-CMV-RasV12), top, or combination with shp53 (pLB-shp53) and PTENi (bottom) and induced with doxycycline. Viability measured using 3D Cell Titer Glo assay and quantified as Relative Luminescence Units (RLU). Data presented is relative to viability at seeding, doxycycline non-induced controls and corrected by values obtained from cultures transduced with respective vector controls (pLENTI-CMV for RasV12, pLB-Scrambled for shp53) or the vehicle control (PTENi). N=3, data are shown as mean ± s.d.; *p < 0.05, **p < 0.01; Student’s *t*-test.
